# PROxAb Shuttle: A non-covalent plug-and-play platform for the rapid generation of tumor-targeting antibody-PROTAC conjugates

**DOI:** 10.1101/2023.09.29.558399

**Authors:** Hendrik Schneider, Sebastian Jäger, Doreen Könning, Nicolas Rasche, Christian Schröter, Desislava Elter, Andreas Evers, Marc Lecomte, Federico Riccardi Sirtori, Daniel Schwarz, Ansgar Wegener, Ingo Hartung, Marcel Rieker

## Abstract

Proteolysis-targeting chimeras (PROTACs) have evolved in recent years from an academic idea to a therapeutic modality with more than 25 active clinical programs. However, achieving oral bioavailability and cell-type specificity remains a challenge, especially for PROTACs recruiting the von Hippel-Lindau (VHL) E3 ligase. Herein, we present an unprecedented, plug-and-play platform for VHL-recruiting PROTACs to overcome these limitations. Our platform allows for the generation of non-covalent antibody-PROTAC complexes within minutes and obviates the need for prior PROTAC modification, antibody-drug linker chemistry optimization or bioconjugation. Our technology relies on camelid-derived antibody domains (VHHs) which can easily be engineered into existing therapeutic antibody scaffolds. The resulting targeted, bispecific fusion proteins can be complexed with PROTACs at defined PROTAC-to-antibody ratios and have been termed PROxAb Shuttles. PROxAb Shuttles can prolong the half-life of PROTACs from hours to days, demonstrate anti-tumor efficacy *in vivo* and have the potential for reloading *in vivo* to further boost efficacy.

## Introduction

Targeted Protein Degradation (TPD) has attracted major interest from both academia and industry over the past years since it enables the selective knockdown of disease-causing proteins with the well-established drug modality of small molecules. The field has been fueled by the more than 25 novel degraders that have advanced into clinical trials within the last 5 years. Degrader modalities have taken various designs, with one of the most established entity being proteolysis-targeting chimeras, or PROTACs, which leverage the ubiquitin-proteasome pathway to degrade and remove disease-causing proteins.^[1,2]^ The inherent advantage of PROTACs lies, among others, in their ability to degrade proteins which have been difficult to address by conventional inhibitors due to the lack of a druggable functional binding pocket. In addition, degrader molecules exhibit a catalytic mode of action that allows strong cellular activity resulting from low compound concentrations. For proteins with slow resynthesis rates, functional effects induced by PROTACs can continue after the PROTAC has already been cleared from the system. The most advanced PROTACs in clinical development are ARV110 and ARV471, which target the androgen and estrogen receptor, respectively and are developed by the company Arvinas.

PROTACs are heterobifunctional molecules that consist of a protein-targeting subunit, a moiety that recruits the respective E3 ligase, and a chemical linker which interconnects both units (Figure 1A). The two E3 ligases most often leveraged for the design of PROTACs are Cereblon (CRBN) and von Hippel-Lindau tumor suppressor (VHL), both belonging to the family of culin ring ligase (CRL) E3 complexes.^[3]^ Although most PROTACs are highly efficient, they are generally not tissue-specific by nature as they exploit VHL and CRBN E3 ligases with broad expression profiles. Tissue-specific degradation of target proteins may be beneficial for minimizing side effects (especially when targeting wild-type proteins), however, functional PROTACs exploiting E3 ligases with restricted tissue distribution are difficult to identify and the development of novel E3 ligase ligands remains a significant challenge.^[4]^ In addition, PROTAC development and efficacy are often hampered by their short half-life *in vivo*^[5,6]^ and low cellular permeability^[7]^, limiting their ability to enter cells and induce protein degradation. The larger size of these chimeric molecules when compared to conventional inhibitors negatively impacts oral bioavailability, solubility, and the absorption, distribution, metabolism and excretion (ADME) properties^[8]^, especially for VHL-targeting PROTACs, due to the peptidic nature of available VHL ligands.^[9]^ Therefore, targeted delivery of PROTACs has evolved as a way to enhance specificity and address the aforementioned liabilities. To this end, covalent conjugation of a PROTAC to an antibody has emerged as an innovative approach that builds on the success of Antibody-Drug Conjugates (ADCs).^[10]^ ADCs rely on the targeted delivery of cytotoxic drugs and have been explored in the clinic for decades, with overall 13 ADCs being approved up to now.^[11]^ The resulting antibody-PROTAC conjugates combine the cellular specificity and favorable pharmacokinetic (PK) profile of the antibody with the protein-degrading ability of the attached PROTAC. While some publications have outlined different strategies for covalent conjugation of the chimeric degrader modality to the antibody, the heterobifunctional nature of PROTACs often impairs the attachment of an additional linker in a suitable exit vector (Figure 1B).^[8,10,12]^ Adding an attachment point to the already bifunctional PROTAC molecule further increases chemical complexity and provides a risk for negatively impacting the biological function of the PROTAC.^[8]^

**Figure 1.**
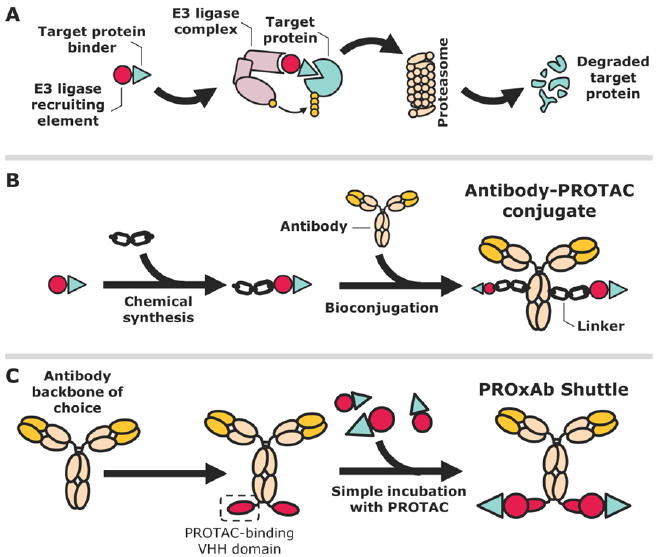
Comparison of the non-covalent PROxAb Shuttle technology with covalent antibody-PROTAC conjugates. A) Mode of action of Proteolysis-targeting chimeras (PROTACs). B) The covalent attachment of a PROTAC to an antibody relies on the identification of a suitable exit vector on the PROTAC for linker attachment. Subsequently, a respective linker-PROTAC molecule needs to be conjugated to the antibody backbone. C) PROxAb Shuttles rely on single domain antibodies derived from camelids which engage the VHL ligand-binding subunit of PROTACs. By fusing such an antibody domain to existing antibody scaffolds, bispecific fusion proteins are obtained.

Recently, alternative TPD modalities that encompass antibodies and membrane-bound E3 ligases have been described. Towards this end, Marei and coworkers generated bispecific, proteolysis-targeting antibodies called PROTABs which bind to a membrane-localized ubiquitin E3 ligase that is selectively expressed in colorectal cancer, as well as to a transmembrane receptor.^[13]^ In addition, the targeted delivery of PROTACs into tumor tissue has also been achieved by fusing the PROTAC to suitable receptor ligands, such as folate. Liu and colleagues could show that fusing a folate moiety to a PROTAC enables the selective delivery of the PROTAC to folate receptor-positive tumor cells.^[14]^ Specific tumor-targeting of PROTACs has also been achieved by chemical conjugation of the degrader to nucleolus-targeting aptamers^[15]^ or amphiphilic diblock copolymers, resulting in self-assembling, micellar nanoparticles.^[16]^ The present work describes a novel TPD platform that involves the non-covalent complexation of PROTACs by bispecific antibodies. This technology leverages single-domain antibodies derived from camelids which bind to the E3 ligase-recruiting subunit of the PROTAC and can be fused to existing therapeutic antibody scaffolds (Figure 1C). The approach relies on the concept of non-covalent interactions between an antibody and a small molecule, similar to previously published approaches by Gupta et al. for the generation of ADCs and several publications authored by scientists at Roche that involve non-covalent binding of small molecules such as biotin or digoxigenin by an antibody.^[17–19]^ Non-covalent approaches have, for instance, been employed for the selective delivery of cytotoxic agents, such as DNA damaging agents, however, the majority of these concepts still relies on the chemical modification of the small molecule with a hapten that is then bound by the antibody.^[20]^ Our platform technology, on the other hand, utilizes the heavy chain-only antibody domains from camelids (termed VHH domains or nanobodies) which directly interact with the VHL-binding portion of the PROTAC.^[21]^ This approach omits the need for the attachment of haptens to the PROTAC and facilitates the generation of non-covalent antibody-PROTAC conjugates.

Camelid-derived single domain antibodies comprise an elongated complimentary determining region 3 (CDR3) and are able to access cryptic or hidden epitopes intractable for conventional antibodies.^[22]^ It has indeed been shown that the elongated CDR3 loop of camelid VHH domains is able to adopt protruding and complex secondary structures that facilitate the interaction with recessed epitopes. As a consequence, VHH domains have been explored for a variety of different targets, including small molecules, and their development culminated in the FDA approval of caplacizumab for the treatment of thrombotic thrombocytopenic purpura (TTP) in 2019.^[23]^ Owing to their small size, VHH domains can also easily be fused to therapeutic antibodies, yielding bispecific and multivalent entities.^[24]^ Our team has generated VHH domains which engage the VHL ligase-recruiting portion of PROTACs. We have fused these domains to the *C*-terminus of therapeutic antibody scaffolds and could show that the resulting fusion proteins, termed PROxAb Shuttles, are able to form stable complexes with PROTACs. Precisely, PROxAb Shuttles non-covalently bind the unmodified PROTAC and the specificity of one VHH allows universal binding to a broad spectrum of different VHL-based PROTACs. Our non-covalent platform enables the rapid complexation of VHL-based PROTACs to antibodies and offers researchers and drug developers a ‘plug-and-play’ approach for the generation of antibody-PROTAC modalities within minutes. Moreover, PROxAb Shuttles are able to drastically prolong the half-life of the PROTAC *in vivo* compared to free PROTAC alone. In addition, PROxAb Shuttles offer the possibility of PROTAC re-loading by re-dosing of the free degrader to further prolong the anti-tumor effect *in vivo*. The technology comes with the benefit of limited synthesis effort as no linker chemistry or bioconjugation techniques are required to generate the resulting PROxAb Shuttle complexes and, as such, the biological function of the degrader is not influenced. The therapeutic antibody of choice needs to be modified only once with the VHL ligand-binding VHH domain and can subsequently be combined with a plethora of different VHL-based degraders in a modular approach. Ultimately, the technology has the potential to improve the therapeutic window of PROTACs that may be too toxic as a standalone therapy, enhance their efficacy by increasing their cell permeability and to improve the *in vivo* PK profile of PROTACs suffering from short half-lives. To the best of our knowledge, this is the first report of non-covalent, PROTAC-binding antibodies to date.

## Results and Discussion

### Anti-VHL ligand antibody domains can be used to convert therapeutic antibody scaffolds into bispecific PROTAC binders (PROxAb Shuttles)

For the generation of anti-VHL ligand antibodies, New World Camelids were immunized with different VH032-haptens (**1**-**4**) conjugated to immunogenic carrier proteins (Figure 2 and Supplementary Information). Of note, VH032 (**5**) is the E3 ligase-engaging structure found in VHL-recruiting PROTACs (Figure 3 and Figure S1). After immunization, titers were determined using enzyme-linked immunosorbent assays (ELISA) and blood samples of camelids were drawn after sufficient titers were achieved. After the isolation of mRNA encoding for the desired VHH domains from peripheral blood mononuclear cells (PBMCs) and translation into cDNA, an antibody phage display library was generated and screened for binding to the VHL-hapten conjugates.

**Figure 2.**
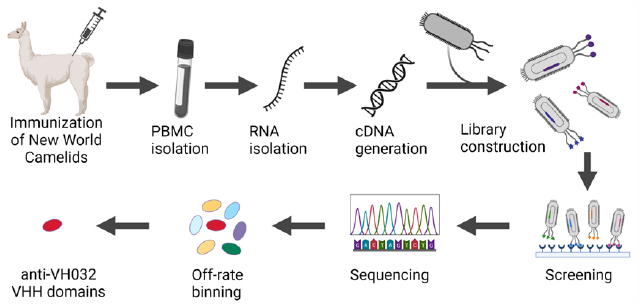
Schematic depiction of the workflow for the generation of VHL ligand-specific antibody domains. Figure created with BioRender.com.

Ultimately, 113 unique antibody clones were identified from the camelid immune library. Prior to reformatting and recombinant expression, monoclonal VHH-displaying phages in bacterial supernatant were subjected to an off-rate screening using Bio-Layer Interferometry (BLI) to identify the most suitable VHH variants for non-covalent delivery with the longest off-rates (Figure S2). Selected VHH domains (termed Monovalent Immobilizer of Chimeras, or MICs) were genetically fused to the *C*-terminus of the antibody scaffold of the therapeutically used anti-CD33 antibody-drug conjugate gemtuzumab ozogamicin (αCD33xMIC) as well as an antibody scaffold that comprised the variable regions of the therapeutic anti-EGFR antibody cetuximab (αEGFRxMIC). The VHH domains and the antibody heavy chains were separated by a short glycine-serine linker. Subsequently, the reformatted fusion proteins were recombinantly expressed in human embryonic kidney (HEK) cells and purified from the cell medium after cellular secretion.

Expression levels and purity of the fusion proteins were comparable to those of the respective antibodies, indicating no negative impact by the fusion of the VHH domains. Next, surface plasmon resonance (SPR) was used to determine the affinity of the VHH-portion of the immobilized fusion proteins towards several VH032-based PROTACs in solution. We selected a broad range of different VH032-based PROTACs comprising aliphatic-, propanediol- and polyethylene glycol (PEG)-based linkers to ensure a certain variability with regard to linker exit vectors and functional groups within the PROTAC (Figure S3). Ultimately, we were able to identify MIC7 as the VHH domain with the highest affinity while accepting a broad diversity of VH032-based PROTACs. As depicted in Tables 1 and S1, MIC7 was able to bind almost all tested PROTACs with linker positions in R1 and R2 with single-digit nano-to subnanomolar affinities (Figure 3).

**Table 1.**
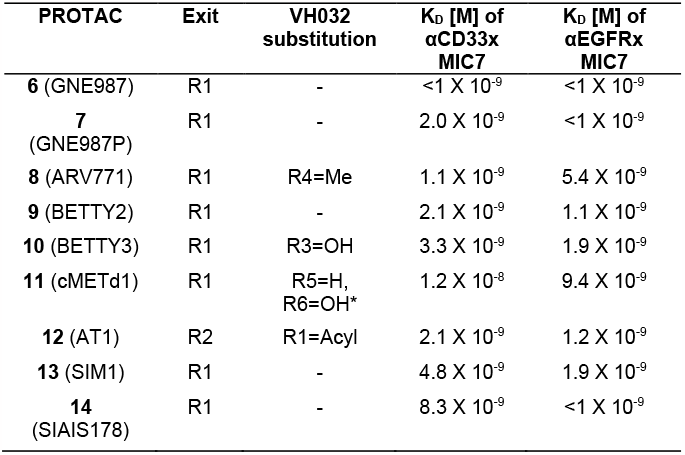
Affinities of reformatted MIC7-based PROxAb Shuttles towards different PROTACs. The MIC7 domain was fused *C*-terminally to the heavy chains of gemtuzumab- and cetuximab-based antibody scaffolds. ^*^ = inactive diastereomer of cMET degrader.

Only one of the tested PROTACs (cMETd1) showed limited binding to MIC7, which was not surprising considering that the hydroxyl group of cMETd1 is present in (*S*)- and not (*R*)-configuration which reduces its ability to engage with VHL and MIC7.

**Figure 3.**
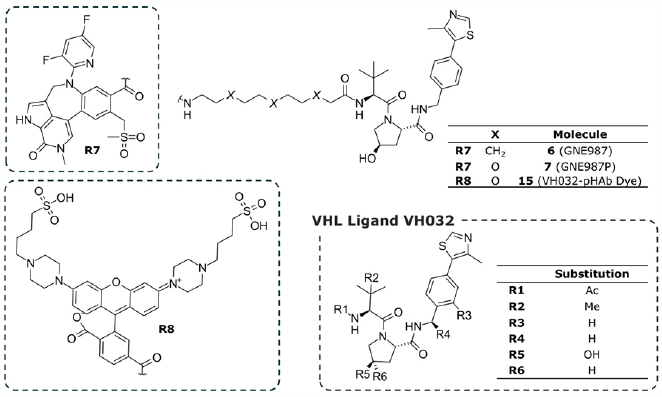
Overview of chemical structures of the PROTACs and tool compounds utilized in the scope of this work.

### Generation of complexed PROxAb Shuttles

We next aimed at investigating the optimal complexation procedure for the generation of PROxAb Shuttles with defined PROTAC-to-antibody ratios. Towards this end, the reformatted, anti-CD33 PROxAb Shuttle comprising the MIC7 VHH domain was incubated with the PROTAC GNE987 (**6**) at room temperature and at a ratio of 2:1 between PROTAC and antibody. GNE987 (**6**) targets the intracellular bromodomain-containing protein 4 (BRD4) and is bound with high affinity by the MIC7 VHH domain (Table 1).^[5]^ The formation of PROxAb Shuttle complexes could already be observed within minutes and successful loading of the PROxAb Shuttle was confirmed by native size exclusion chromatography (SEC) coupled with native mass spectrometry (MS). As depicted in Figure 4A, the complexed and non-complexed αCD33xMIC7 Shuttles with a PROTAC-to-antibody ratio of 1.9 and 0, respectively, could be identified. Upon using native SEC coupled with native MS, it was possible to verify the loading of PROxAb Shuttles with defined PROTAC-to-antibody ratios. In addition, isothermal titration calorimetry (ITC) was used to determine the stoichiometry of the antibody/VH032 interaction, as depicted in Figure 4B which shows the binding isotherm for αCD33xMIC7 complexed with VH032. In this experiment, we observed symmetrical isotherms and interactions which were dominated by binding enthalpy. The binding is saturating at an apparent molar ratio of ∼0.5, which is equivalent to a stoichiometry of N = 2.1. These results are in line with the expected ratio of two VH032 molecules binding to one PROxAb Shuttle comprising two MIC7 VHH entities. The symmetrical isotherm depicted in Figure 4B indicates a homogeneous binding process with one set of identical and independent binding sites.

**Figure 4.**
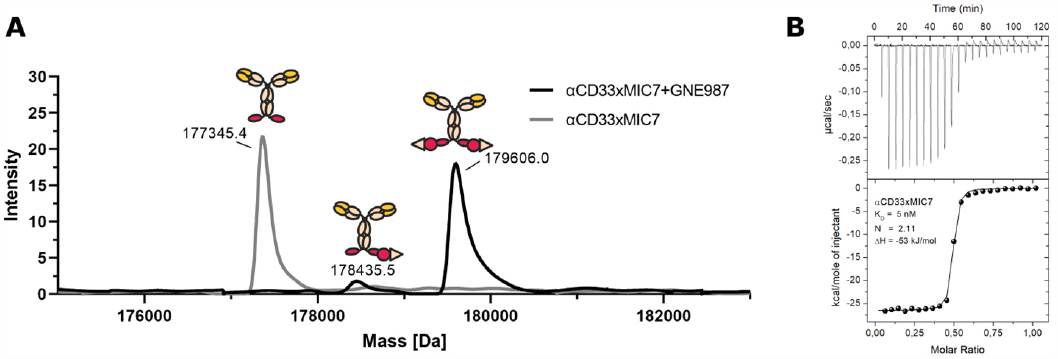
Complexation process and stoichiometry of the antibody/PROTAC interaction. A) Spectrum of native SEC/native MS measurements for the αCD33xMIC7 PROxAb Shuttle complexation reaction with the PROTAC GNE987 (**6**). Distinct peaks for the PROxAb Shuttles with a PROTAC-to-antibody ratio of 0 and 2 can be identified, respectively. B) Binding isotherm of the αCD33xMIC7 PROxAb Shuttle with the VHL ligand VH032 (**5**).

### PROxAb Shuttles efficiently internalize into cells and exhibit nanomolar potencies *in vitro*

To assess whether the cellular binding of our PROxAb Shuttles was impaired by the fusion of the MIC7 VHH domain, cell lines with varying CD33 receptor expression were incubated with either unmodified antibody, PROTAC-complexed αCD33xMIC7 PROxAb Shuttle, or an isotype control for 45 minutes. An antibody targeting digoxigenin, which was likewise fused to MIC7 at its *C*-terminus, served as isotype control (αDIGxMIC7). Following the incubation, the cells were stained with a fluorescently labeled anti-human Fc detection antibody and subjected to flow cytometric analyses. The resulting mean fluorescence intensity (MFI) values for the MIC7-based PROxAb Shuttles, complexed or non-complexed with GNE987 (**6**), were compared to the parental αCD33 antibody and are depicted in Figure 5A, demonstrating that the fusion of the MIC7 VHH domain did not impair binding to CD33.

**Figure 5.**
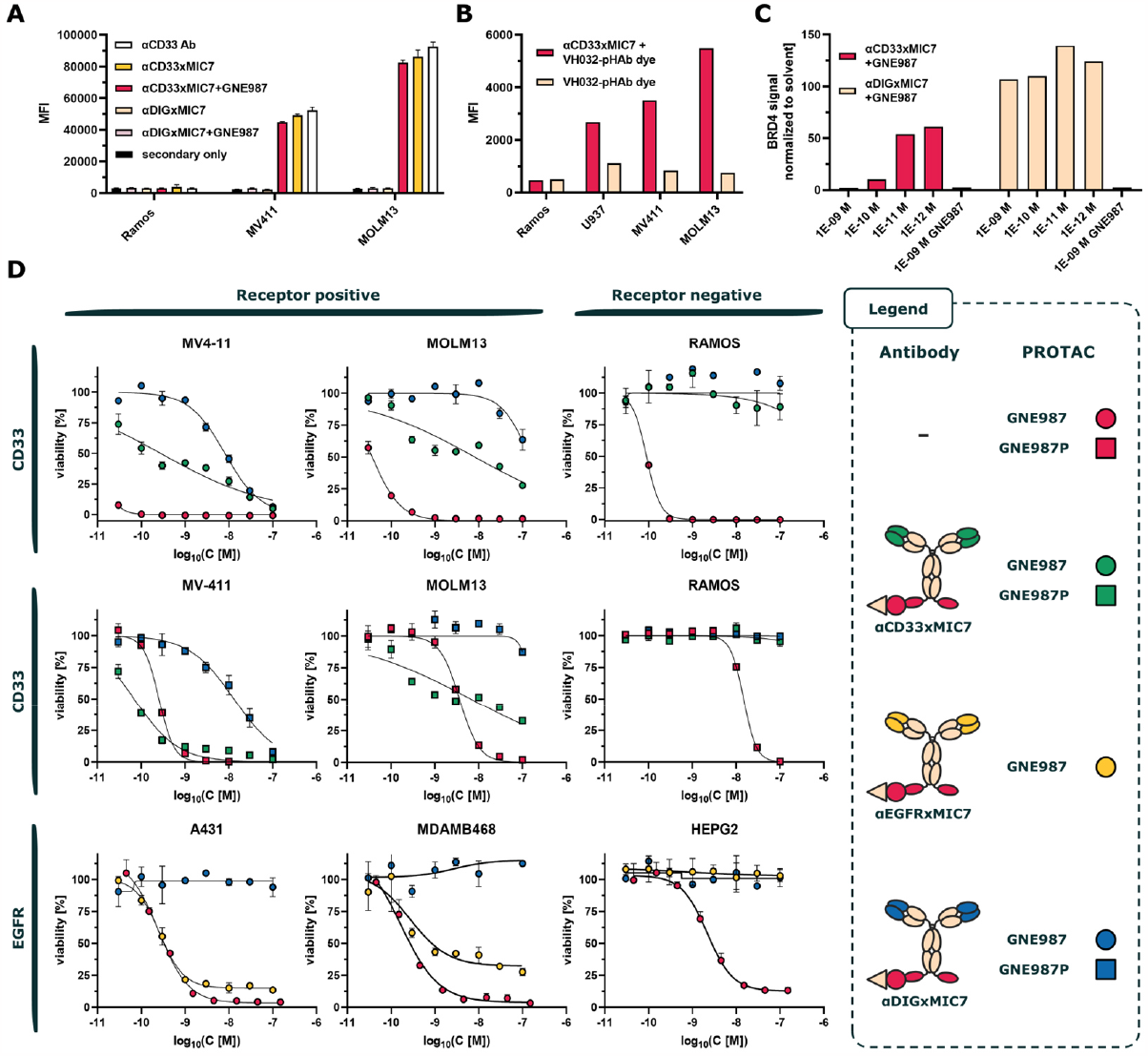
*In vitro* characterization of MIC7-based PROxAb Shuttles. A) Cell binding of anti-CD33 antibody scaffolds with and without a *C*-terminally fused MIC7 antibody domain to CD33 positive cells. PROxAb Shuttles were tested in the complexed (PROTAC-to-antibody ratio = 1) and non-complexed form with GNE987 PROTAC (**6**). Bars represent the average of technical duplicates. Ab = antibody. B) Flow cytometric analysis of the internalization of the αCD33xMIC7 PROxAb Shuttle complexed with VH032-pHAb dye (**15**) over a period of 24 hours (dye-to-antibody ratio = 1) compared to free dye alone on different CD33-expressing cell lines. C) Western Blot analysis and quantification of BRD4 levels in CD33-positive cells treated with different concentrations of αCD33xMIC7 PROxAb Shuttle complexed with GNE987 or free GNE987 (**6**) at a PROTAC-to-antibody ratio of 1. The intensity of individual Western Blot signals was normalized to the loading control (Actin; Figure S4). D) Cell viability assays of complexed αCD33- and αEGFRxMIC7 PROxAb Shuttles with different PROTACs on target-positive and target-negative cell lines at PROTAC-to-antibody ratios of 1. Compounds were incubated on cells for overall 72 hours. An αDIGxMIC7 Shuttle was included as an isotype control and free PROTAC was used as the positive control for each assay. Upper panel: Dose-response curves of αCD33xMIC7 Shuttles and isotype control complexed with GNE987(**6**). Middle panel: Dose-response curves of αCD33xMIC7 PROxAb Shuttle and isotype control complexed with GNE987P (**7**). Lower panel: Dose-response curves of αEGFRxMIC7 PROxAb Shuttle and isotype control complexed with GNE987 (**6**). Sigmoidal curves represent the mean of technical duplicates.

To gain further insights into the route of internalization of the PROxAb Shuttle, we synthesized a VH032 subunit that was linked to a pH-responsive dye (VH032-pHAb dye (**15**); Figure 3) via a PEG-based linker. The shift in environmental pH that occurs upon transition of the PROxAb Shuttle-bound VH032-pHAb dye (**15**) into the acidic environment of the endosomal and lysosomal vesicles causes an increase in the fluorescence signal. After incubation of CD33-positive cells with VH032-pHAb-complexed αCD33xMIC7 PROxAb Shuttle, the cells were subjected to flow cytometric analyses. As depicted in Figure 5B, the MFI values increased when cells were incubated with the PROxAb Shuttle complex in comparison to the corresponding VH032-pHAb dye-only control (**15**), demonstrating internalization of the Shuttle and accumulation of the VH032-pHAb dye (**15**) in the endolysosomal vesicles.

In order to show that PROTAC which is internalized via the PROxAb Shuttle also reaches its intracellular target, we next measured the degradation of BRD4 after incubation of CD33-positive MV4-11 cells with decreasing concentrations of GNE987-complexed αCD33xMIC7 and αDIGxMIC7 PROxAb Shuttles or GNE987 PROTAC (**6**) alone (Figure 5C). Following the incubation, cells were lysed at different time points and the cell lysates were subjected to Western Blot analyses to determine the remaining levels of BRD4 (Figure S4). Figure 5C shows the quantified BRD4 signal from the Western Blots in % and normalized to the solvent-only control. Upon incubation of the cells with the PROxAb Shuttles, a concentration-dependent decrease of the intracellular BRD4 signal could be observed for the αCD33xMIC7 Shuttle and for free PROTAC, whereas BRD4 signals were not impacted by the isotype control. These results demonstrate that the PROTAC, when delivered with a PROxAb Shuttle, can escape the lysosomal compartment to exert its intended mode of action. Moreover, these data demonstrate the stability of the PROxAb Shuttle complex under the assay conditions since the isotype control did not elicit BRD4 degradation. Notably, complete BRD4 degradation was achieved by the αCD33xMIC7 Shuttle at a concentration as low as 1 nM. Ultimately, we investigated whether it is possible to deliver PROTAC concentrations into tumor cells via the PROxAb Shuttle that are sufficient to elicit the intended pharmacological effect of cell killing. Since the PROxAb Shuttle complexes can be freshly prepared prior to each potency assay without any further purification step, we first performed titration experiments with different PROTAC-to-antibody ratios on target-negative cell lines (data not shown). This served to determine the ratio which would yield the lowest background signal to enable an accurate assessment of the median inhibitory concentrations of our PROxAb Shuttles in subsequent experiments. Ultimately, a PROTAC-to-antibody ratio of 1 was selected for the *in vitro* potency assays. Finally, CD33- and EGFR-targeting PROxAb Shuttles were incubated with either, CD33- or EGFR-positive and negative tumor cell lines with varying receptor expression. The αDIGxMIC7 PROxAb Shuttle served as isotype control and free PROTAC was included as a positive control. Figure 5D depicts the resulting dose-response curves for each individual cell line and for each PROxAb Shuttle with a PROTAC-to-antibody ratio of 1. The complexed αCD33xMIC7 and αEGFRxMIC7 Shuttles exhibited potencies on target positive cells in the nanomolar range and were able to selectively deliver cytotoxic concentrations of PROTACs to the desired cells (Table S2).

Notably, the cytotoxicity of each Shuttle was also dependent on the number of target receptors on the cell surface. Whereas the EGFR copy numbers are high and in the range of several hundred thousand for the tested A431 and MDA-MB-468 cells,^[25]^ the CD33-positive cell lines MV4-11 and MOLM13 comprise a lower receptor number of around 18,000 to 48,000 copies.^[26,27]^ These differences are reflected in the higher half maximal inhibitory concentration (IC_50_) values of the GNE987-complexed αCD33xMIC7 Shuttle when compared to free PROTAC alone (Figure 5D, upper panel), while the complexed aEGFRxMIC7 Shuttle shows a similar potency in comparison to the free PROTAC. In addition to GNE987 (**6**), we also complexed the αCD33xMIC7 Shuttle with the more hydrophilic variant of GNE987(**6**), termed GNE987P (**7**).^[12]^ Combining a more hydrophilic and, therefore, less cell permeable PROTAC such as GNE987P (**7**) with the PROxAb Shuttle technology should enable to boost PROTAC potency due to a facilitated active delivery into the target cell. As can be seen in the middle panel of Figure 5B and in Table S2, GNE987P (**7**) in complex with our αCD33xMIC7 Shuttle was more efficacious when compared to the free PROTAC alone, particularly on MV4-11 cells. Importantly, no cytotoxic effects were observed for all targeted PROxAb Shuttles on either, the CD33- or EGFR-negative cell lines regardless of the complexed PROTAC. After these initial experiments, we were able to demonstrate similar effects for numerous PROxAb Shuttles based on different therapeutic antibody scaffolds, also in combination with other PROTACs than GNE987 (**6**) and GNE987P (**7**), which further emphasizes the platform potential of PROxAb Shuttles (Table S3).

### PROxAb Shuttles can extend the half-life of PROTACs from hours to days

After demonstrating that PROxAb Shuttles effectively deliver VHL-based PROTACs *in vitro*, we next investigated their pharmacokinetic (PK) profile in C57BL/6N mice. In order to show that the overall PK of the antibodies was not altered by the VHH fusion, we first compared the PK parameters of the unmodified αCD33 antibody with 2 different αCD33 PROxAb Shuttles without complexed PROTAC (Figure 6A). Besides the MIC7 Shuttle, a second αCD33 PROxAb Shuttle comprising the VHL-binding VHH domain MIC5 was included as well (αCD33xMIC5) in order to assess the influence of different antibody off-rates on the pharmacokinetics of the complexed PROTAC in further experiments. MIC5 exhibits a lower affinity towards VH032-based PROTACs as well as a faster off-rate when compared to MIC7, as assessed in previous surface plasmon resonance experiments (Table S4). Table 2 summarizes the pharmacokinetic parameters for the respective antibodies at an administered dose of 30 mg/kg. All tested antibody variants exhibited half-lives of approximately 10 days, emphasizing that the PK profile of the αCD33 antibody is not influenced by the *C*-terminal fusion of the MIC5 and MIC7 VHH domains. Moving forward, we evaluated the αCD33xMIC7 and αCD33xMIC5 Shuttles with and without bound PROTAC in order to confirm that the associated PROTAC does not influence the PK property of the fusion protein. In contrast to previous *in vitro* assays, the PROTAC-to-antibody ratio was set to 2 for all *in vivo* experiments to reach a higher complexed PROTAC exposure and, therefore, to increase the efficacy of the Shuttles. Figure 6B shows that the total antibody concentration profile in plasma as assessed *via* electrochemiluminescence immunoassay, is nearly identical for the complexed and non-complexed αCD33 PROxAb Shuttles, indicating a similar PK profile for the antibody.

**Table 2.** Pharmacokinetic parameters for different PROTACs, antibodies and either complexed or non-complexed PROxAb Shuttles with PROTAC-to-antibody ratios of 2. Abbreviations: t_1/2_ = half-life time; Cl = clearance, Vss = volume of distribution at steady-state; TAb = total antibody.

| Molecule | Analyte | $t_{1/2}$ [h] | $C_{max}$ [ng/mL] | $AUC_{0-inf}$ [h x ng/mL] | Cl [mL/h/kg] | Vss [mL/kg] |
| --- | --- | --- | --- | --- | --- | --- |
| 6 (GNE987) | PROTAC | 2.2 | $7.2 \times 10^2$ | $3.5 \times 10^2$ | 850 | 1348 |
| 7 (GNE987P) | PROTAC | 0.54 | $2.8 \times 10^3$ | $1.3 \times 10^3$ | 236 | 86 |
| αCD33 antibody | TAB | 237 | $8.1 \times 10^5$ | $2.2 \times 10^8$ | 0.14 | 74 |
| αCD33xMIC5 | TAB | 351 | $6.6 \times 10^5$ | $1.7 \times 10^8$ | 0.22 | 85 |
| αCD33xMIC5 with 6 | PROTAC | 30 | $7.9 \times 10^3$ | $1.3 \times 10^5$ | 3.0 | 100 |
| αCD33xMIC5 with 6 | TAB | 276 | $7.2 \times 10^5$ | $1.7 \times 10^8$ | 0.18 | 67 |
| αCD33xMIC7 | TAB | 274 | $6.8 \times 10^5$ | $1.6 \times 10^8$ | 0.19 | 75 |
| αCD33xMIC7 with 6 | PROTAC | 105 | $2.5 \times 10^3$ | $2.3 \times 10^5$ | 1.74 | 270 |
| αCD33xMIC7 with 6 | TAB | 304 | $5.9 \times 10^5$ | $1.3 \times 10^8$ | 0.25 | 101 |
| αCD33xMIC7 with 7 | PROTAC | 34 | $5.6 \times 10^3$ | $1.0 \times 10^5$ | 3.8 | 140 |
| αEGFRxMIC7 with 6 | PROTAC | 66 | $3.2 \times 10^3$ | $1.3 \times 10^5$ | 3.0 | 280 |

**Figure 6.**
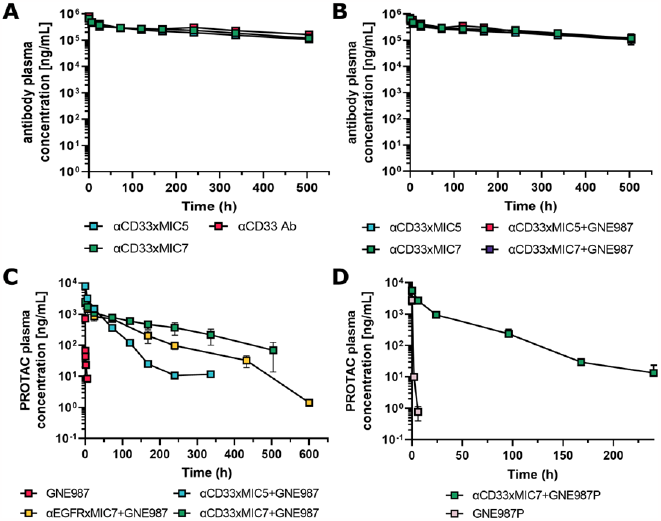
Pharmacokinetic profiles of complexed (PROTAC-to-antibody ratio of 2) and non-complexed PROxAb Shuttles (30 mg/kg; i.v. route) with GNE987 (**6**) (0.3 mg/kg) in mice. A) Comparison of the total antibody concentration profiles of unmodified αCD33 antibody scaffold with non-complexed αCD33xMIC7 and αCD33xMIC5 PROxAb Shuttles.B) Total antibody concentration profiles of complexed and non-complexed αCD33xMIC7 and αCD33xMIC5 PROxAb Shuttle variants. C) Total PROTAC profiles of αCD33xMIC7, αCD33xMIC5 and αEGFRxMIC7 Shuttles after complexation with GNE987 (**6**), as well as free GNE987 (**6**). D) Total PROTAC profiles of the αCD33xMIC7 PROxAb Shuttle after complexation with GNE987P (**7**), as well as free GNE987P (**7**).

Next, we investigated the influence of the PROxAb Shuttle on the half-life of the PROTAC. To achieve this, C57BL/6N mice were dosed with either 30 mg/kg of GNE987-complexed αCD33xMIC7 and αCD33xMIC5 Shuttles (comprising overall 0.38 mg/kg of GNE987 **(6)**) or a dose of 0.3 mg/kg of GNE987 alone (**6**). As can be seen in Figure 6C, the half-life of GNE987 (**6**) dosed alone was approximately 2 hours and could be drastically prolonged to more than 4 days (105 hours) when complexed by the αCD33xMIC7 Shuttle. Interestingly, the half-life of GNE987 (**6**) when complexed by the MIC5-based PROxAb Shuttle could solely be increased up to 30 hours, indicating that affinity is playing a role for efficient and long-lasting capture of the PROTAC *in vivo*. The difference in affinity between MIC5 and MIC7 for GNE987 (**6**), as measured via surface plasmon resonance, was almost 17-fold and translated into a 3.5-fold higher PROTAC half-life in mice. Notably, the off-rates (k_off_) of both antibody domains varied by a factor of 21, while the on-rates (k_on_) were in the same range (Table S4). Fine-tuning of the antibody/PROTAC affinity by, for instance, affinity maturation of MIC7 with a focus on dedicated off-rate screenings may result in even tighter interactions and even longer PROTAC *in vivo* half-lives. It is worth noticing that the MIC5 and the MIC7-based αCD33 PROxAb Shuttle complexes were well-tolerated in treated mice, and no loss of body weight or any clinical signs of toxicity were observed (data not shown).

Building on these findings, we next investigated the PK properties of the αEGFRxMIC7 Shuttle in order to extend our previous analyses to another antibody scaffolds. Similar to the results obtained for the αCD33 Shuttle variants, the αEGFRxMIC7 PROxAb Shuttle was able to prolong the *in vivo* half-life of GNE987 (**6**) from 2 hours to approximately 66 hours as compared to the PROTAC alone (Figure 6C).

To further demonstrate the versatility of the PROxAb Shuttle technology with regards to the PROTAC, we ultimately thought to investigate the combination of αCD33xMIC7 PROxAb Shuttles with another PROTAC than GNE987 (**6**). For this purpose, the αCD33xMIC7 Shuttle was complexed with GNE987P (**7**) at a PROTAC-to-antibody ratio of 2 and administered at 30 mg/kg (comprising overall 0.38 mg/kg of GNE987P (7)) while a second cohort was dosed with 0.3 mg/kg of GNE98P alone (**7**) (Figure 6D). In this case, the half-life of GNE987P (**7**) could be increased from 0.5 to 33.6 hours, i.e. a factor of >60-fold. These results further validate the *in vivo* half-life-prolongation effect of the PROTAC that is achievable by the PROxAb Shuttle concept.

Leveraging a non-covalent rather than a covalent bond between the antibody and the PROTAC has the advantage of potential re-binding to the antibody. This means, that PROTACs which may be released from the Shuttle in circulation can in fact be re-bound by the antibody. Fine-tuning of the antibody/PROTAC interaction by additional protein engineering may result in tailor-made ‘PROTAC reservoirs’ that could balance out suboptimal plasma stability by slow release from the Shuttle.^[28]^ In fact, multiple non-covalent interactions can even surpass a single covalent bond with regards to affinity and form an almost irreversible connection.^[17]^

### PROTAC-complexed αCD33xMIC7 PROxAb Shuttles elicit antitumor responses *in vivo*

To demonstrate that the PROxAb Shuttle technology not only leads to a drastic half-life prolongation but also enables an efficient delivery of PROTACs into tumors, mouse xenograft studies have been performed. Since the αCD33xMIC7 PROxAb Shuttle complexed with GNE987 demonstrated the highest PROTAC half-life prolongation in the previous PK studies, this combination was further investigated regarding its potential to slow down tumor growth in a tumor-bearing mouse model. The αCD33xMIC7 PROxAb Shuttle was complexed with GNE987 (**6**), the most potent PROTAC in our *in vitro* assays (Tables S1 and S2), at a PROTAC-to-antibody ratio of 2 and the myelomonocytic leukemia cell line MV4-11 was selected as a suitable *in vivo* model. MV4-11 has a moderate CD33 receptor density and was shown to be sensitive towards both, free GNE987 (**6**) and the respective complexed PROxAb Shuttles in previous *in vitro* experiments (Figure 5D). The administered doses of complexed PROxAb Shuttle were matched to equivalent amounts of PROTAC in order to enable a direct comparison between the mice treated with complexed PROxAb Shuttle and PROTAC alone. Figure 7A depicts the tumor growth inhibition curves in female CB17 SCID mice treated either with GNE987 alone (**6**) or complexed and non-complexed αCD33xMIC7 Shuttle, or complexed αCD33xMIC5 Shuttle and a vehicle control. All tested molecules were administered intravenously on day 1 as a single dose. The complexed αCD33xMIC7 PROxAb Shuttle demonstrated superior antitumor activity as compared to an equivalent dose of GNE987 (**6**) alone or the vehicle control (Figure 7C). In addition, the respective non-complexed αCD33xMIC7 Shuttle did not elicit any antitumor effect at all. These results are in line with the prolonged half-life of the PROTAC when complexed with a PROxAb Shuttle, as observed during previous PK studies, and indicate that targeted tumor drug delivery of a PROTAC via the non-covalent PROxAb Shuttle approach results in similar effects as observed for conventional ADCs.^[29]^ Also, the αCD33xMIC5 Shuttle complexed with GNE987 (**6**) was not able to induce a significant antitumor response when compared with the vehicle control, which can likely be attributed to the 21-fold faster off-rate of the MIC5 antibody domain (Table S4). In addition to the accelerated complexation and generation of PROxAb Shuttles, another advantage of the non-covalent versus the covalent interaction could be the application of therapeutic pre-targeting concepts wherein the targeting antibody and the PROTAC are administered separately to enable binding of the antibody to the cells of interest in a first step. Especially for toxic small molecules, pre-targeting has the advantage of minimizing systemic exposure and, therefore, potentially unwanted side effects associated with such non-targeted molecules.^[30,31]^ Recently, Bordeau and coworkers described an approach for a non-covalent and therapeutically focused antibody/small molecule interaction which is based on the concept of re-binding *in vivo*.^[32]^ Their work focused on an antibody that was able to bind to the cytotoxic payload monomethyl auristatin E (MMAE), a toxin that is frequently used in ADC design. The approach aimed at the detoxification of free MMAE originating from the degradation of an MMAE-based ADC *in vivo*, and depends on the co-administration of their anti-MMAE antibody with the ADC. Bordeau and coworkers could demonstrate that the anti-MMAE antibody and free, deconjugated MMAE could bind to one another *in vivo* and, consequently, could diminish common MMAE-related side effects. Intrigued by this concept, we next wanted to elucidate the possibility of re-binding for the PROxAb Shuttle technology. Prior to conducting efficacy experiments in tumor-bearing mice, we first investigated the re-binding ability of a PROxAb Shuttle by performing PK studies in which we sequentially administered a non-complexed αCD33xMIC7 PROxAb Shuttle followed by administration of the PROTAC GNE987P (**7**) 24 hours later (Figure S5). The resulting plasma profiles of pre-complexed αCD33xMIC7 were nearly identical to the plasma profile after sequential administration of the αCD33xMIC7 Shuttle and GNE987P (**7**) (with half-lives of 33.6 and 25.2 hours, respectively), demonstrating the possibility of *in vivo* complexation between our PROxAb Shuttle and free PROTAC. Motivated by these results, we treated MV4-11 tumor-bearing mice with the αCD33xMIC7 PROxAb Shuttle complexed with GNE987 (**6**) on day 1 (PROTAC-to-antibody ratio of 2) and re-dosed mice with GNE987 (**6**) alone on day 8. This interval was chosen based on previous total antibody plasma profiles of αCD33xMIC7. Of note, the dose of GNE987 (**6**) administered on day 8 was equivalent to the dose received on day 1. As can be seen in Figure 7C and E, re-dosing of the PROTAC to the Shuttle could prolong the efficacy of the αCD33xMIC7 PROxAb Shuttle by several days as compared to the group that did not undergo the re-dosing treatment. Moreover, the antitumor effect of the PROxAb Shuttle was significantly more pronounced when compared to the group that solely received doses of free PROTAC on day 1 and day 8. Importantly, the PROxAb Shuttles were well-tolerated, and no body weight loss was observed (Figure S6). Dedicated experiments that focus on sequential administration of a non-complexed Shuttle and PROTAC alone are currently under investigation by our group. Such studies could open avenues for the direct assembly of non-covalent ADCs *in vivo* and for utilizing sequential loading of one PROxAb Shuttle with PROTACs targeting different proteins to maximize patient benefit in a highly personalized approach.

**Figure 7.**
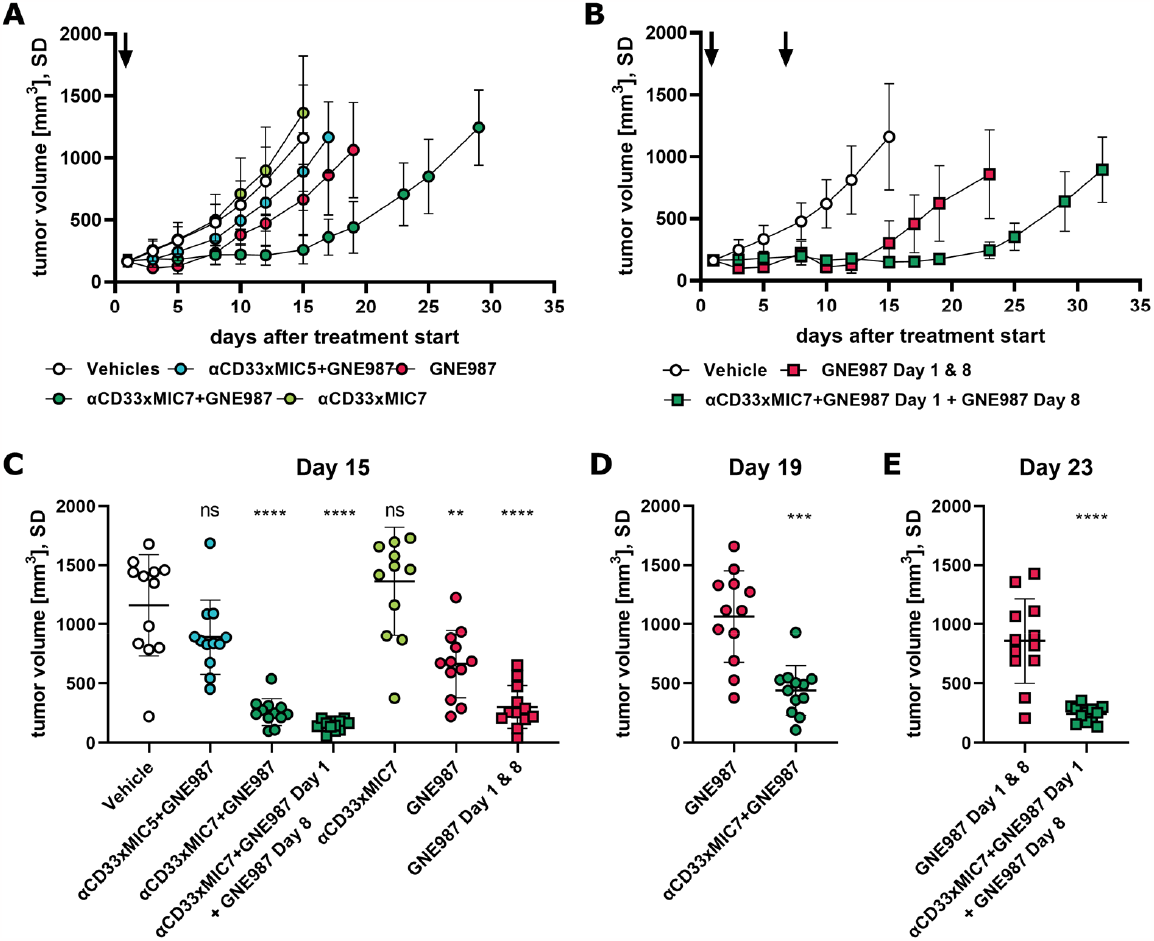
Results of *in vivo* efficacy studies in tumor-bearing mice. A) The αCD33xMIC5 and αCD33xMIC7 PROxAb Shuttles in a complexed and non-complexed form were administered to MV4-11 tumor-bearing mice at a dose of 30 mg/kg. The PROTAC-to-antibody ratio was set to 2 for each Shuttle. Free GNE987 PROTAC (**6**) was administered at a dose equivalent to complexed GNE987 (**6**) in the PROxAb Shuttles (i.e. 0.38 mg/kg). The arrow indicates the time of drug administration. B) Free GNE987 (**6**) as well as GNE987-complexed αCD33xMIC7 PROxAb Shuttle were administered to MV4-11 tumor-bearing mice at doses of 0.38 mg/kg and 30 mg/kg, respectively. Re-dosing of GNE987 (**6**) at 0.38 mg/kg was performed in both groups on day 8. Arrows indicate the time of drug administration. C), D) and E) End point analyses of the *in vivo* efficacy study depicted in A) on days 15, 19 and 23, respectively. Significance levels were determined by Two-Way ANOVA (C) and two-tailed, unpaired t-Test (D and E).

### The camelid-derived MIC7 VHH antibody domain can be humanized to yield therapeutic PROxAb Shuttle candidates

The reduction of potential immunogenicity and the associated side effects, such as rapid clearance or neutralization of the antibody drug, remain an essential pillar in antibody drug development. To address this, we sought to humanize the camelid-derived antibody domain MIC7 which we employed in our best-performing PROxAb Shuttles. Humanization of camelid antibody domains was first described by Vincke and coworkers in 2009 and was since developed into an established protocol that is widely applied to therapeutic nanobodies, culminating in the approval of the humanized nanobody caplacizumab.^[33,34]^ Leveraging our recently described approach for *in silico* sequence assessment, the humanization strategy that was employed for MIC7 started with a sequence-alignment with the most similar human germline sequence and, subsequently, relied on the modification of residues which were located outside of critical structural positions (Vernier and Hallmark positions) in order to preserve the structural integrity of the CDRs.^[35]^ Building on these initial mutations, additional humanized MIC7 variants were generated by adding the humanized residues in these key structural positions in a step-by-step process. The resulting humanized MIC7 mutant domains (hMIC7_1.X) were fused to the αCD33 antibody to generate full-length PROxAb Shuttles. The affinity towards VH032 (**5**) was confirmed *via* ITC (Table S5) and SPR (Table S6).

Using ITC, we could also confirm the stoichiometries of N = 2 or 2.1 for all humanized MIC7 variants. The fully humanized αCD33xhMIC7_1.6 Shuttle variant comprising the most mutations was able to elicit selective cell killing in the same potency range as αCD33 PROxAb Shuttles with humanized MIC7 domains (Table 3). Taken together, hMIC7_1.6-based PROxAb Shuttles performed similar when compared to their non-humanized MIC7 precursor in terms of protein production, stability, binding affinity as well as *in vitro* cytotoxicity.

**Table 3.** Potencies of humanized and wildtype αCD33 PROxAb Shuttles complexed with GNE987 (**6**) or GNE987P (**7**) at a PROTAC-to-antibody ratio of 1 on CD33-positive MV4-11 and MOLM13 as well as CD33-negative RAMOS cells. Potencies of PROTAC GNE987 (**6**) and GNE987P (**6**) alone have also been included for comparison.

| Shuttle | PROTAC | IC<sub>50</sub> [M] |  |  |
| --- | --- | --- | --- | --- |
|  |  | MV4-11 | MOLM13 | RAMOS |
| αCD33xhMIC7_1.6 | 6 | <3.0 × 10<sup>-11</sup> | 6.6 × 10<sup>-11</sup> | > 1.0 × 10<sup>-7</sup> |
| αCD33xMIC7 | 6 | 3.6 × 10<sup>-11</sup> | 1.3 × 10<sup>-9</sup> | > 1.0 × 10<sup>-7</sup> |
| - | 6 | 6.7 × 10<sup>-12</sup> | 2.3 × 10<sup>-11</sup> | 1.3 × 10<sup>-10</sup> |
| αCD33xhMIC7_1.6 | 7 | 7.9 × 10<sup>-11</sup> | 9.2 × 10<sup>-10</sup> | > 1.0 × 10<sup>-7</sup> |
| αCD33xMIC7 | 7 | 8.4 × 10<sup>-11</sup> | 2.1 × 10<sup>-9</sup> | > 1.0 × 10<sup>-7</sup> |
| - | 7 | 2.9 × 10<sup>-10</sup> | 1.6 × 10<sup>-9</sup> | 1.7 × 10<sup>-8</sup> |

## Conclusion

Rising interest in the field of Targeted Protein Degradation has culminated in the transition of the first PROTACs into advanced clinical trials.^[1]^ Despite their catalytic mode of action and ability to engage therapeutic targets which are often intractable for conventional inhibitors, developing cell type-specific PROTACs remains a challenge. Covalent antibody-PROTAC conjugates have recently been reported, combining the cellular specificity of an antibody with the cell killing properties of a PROTAC.^[4,5,8,12]^ Nevertheless, the heterobifunctional nature of PROTACs often impedes the identification of suitable and generic attachment sites for chemical linkers, resulting in time- and resource-consuming efforts. Herein, we describe the identification and therapeutic application of camelid-derived single domain antibodies which bind to the E3 ligase-recruiting subunit of a PROTAC with single-digit nano-to subnanomolar affinities. Specifically, antibodies that engage VH032-based PROTACs which recruit the VHL E3 ubiquitin ligase, one of the most frequently leveraged ligases for PROTAC design, have been identified in the scope of this work. Fusing the camelid antibody domain to therapeutic human antibody scaffolds resulted in bispecific, multivalent fusion proteins that build strong non-covalent complexes with PROTACs at defined PROTAC-to-antibody ratios. These complexes effectively deliver PROTACs to target cells and selectively induce cell killing. Notably, the *in vivo* half-life of PROTACs bound by PROxAb Shuttles was prolonged from a few hours to several days and the complexes induced antitumor responses in mouse xenograft models comparable to effects observed for conventional ADCs. In addition, we demonstrated that re-dosing of the PROTAC alone to tumor-bearing mice which were pre-dosed with complexed PROxAb Shuttle can further prolong tumor growth control by the Shuttle. Fine-tuning of PK- and efficacy parameters by affinity maturation of the antibody/PROTAC interaction is expected to enable the design of PROxAb Shuttles with tailor-made characteristics. The therapeutic potential of this approach is further underlined by the straightforward humanization strategies that have been implemented for camelid-derived VHH domains over the past years and that have culminated in the FDA-approval of the humanized VHH variant caplazicumab.^[23,33,34]^

Our non-covalent PROxAb Shuttle complexes can be generated rapidly and in a matter of only a few minutes at the bench whereas the covalent conjugation of PROTACs to antibodies has proven to be challenging and time-consuming in the past. The PROxAb Shuttle technology could enable scientists to assess degrader candidates which have not been optimized with regards to PK profile or cellular permeability, both *in vitro* and *in vivo*. In summary, the PROxAb Shuttle technology represents a plug-and-play platform for the rapid generation of VH032-based antibody-PROTAC complexes and has the potential to rescue PROTACs which may not be suitable for clinical development as a monotherapy due to limitations regarding *in vivo* toxicity or reaching sufficient oral bioavailability.

## Acknowledgements

The authors would like to sincerely thank the following people at Merck KGaA, Darmstadt, Germany who have contributed to the experiments and results presented in the scope of this publication: Andreas Schönemann for SPR measurements, Frank Fischer, Djordje Musil, Martin Lehmann, Ralf Günther, Thomas Eichhorn, Sabine Raab-Westphal, Kevin Heitland, Stephanie Czasch, Ilse de Salve, Elisa Bertotti, Patrizia Tavano, Jens Hannewald, Ingrid Schmidt, Laura Basset, Eva Maria Leibrock, Yvonne Bischoff, Vanessa Lautenbach and Anja Ogroske. In addition, the authors would like to thank Paul Gehrtz for the synthesis of the VH032-pHAb dye and the following companies for their services: preclinics GmbH, YUMAB GmbH and Reaction Biology Corp.

