## supplemental_information for "PROxAb Shuttle: A non-covalent plug-and-play platform for the rapid generation of tumor-targeting antibody-PROTAC conjugates": PROxAb Shuttle A non-covalent plug-and-play platform for the rapid generation of tumor-targeting antibody-PROTAC conjugates supplemen.docx

**Supporting Information**

### **Experimental Procedures**

#### Materials

All chemicals, building blocks and reagents were purchased from commercial sources such as Sigma-Aldrich Corp., abcr GmbH, Thermo Fisher Scientific Inc., Merck KGaA, Bio-Techne GmbH, and VWR International, Medchemexpress®, and used without any further purification unless otherwise stated.

(S,R,S)-AHPC-PEG_3_-NH_2_ hydrochloride, and (S,R,S)-AHPC-C_6_-CO_2_H hydrochloride **4** (VHL-c) were acquired from Sigma Aldrich. GNE987 (**6**), GNE987P (**7**, PROTAC BRD4 Degrader-8), ARV771 (**8**), SIM1 (**13**), SIAIS178 and (**14**) were acquired from MedChemExpress®, USA. AT1 (**12**) was acquired from Tocris Biosience, UK. All PROTACs and building blocks were used without any further purification unless otherwise stated.

##### Synthesis of Hapten 1 (VHL-1)

2-(2-(2-(((S)-1-((2S,4R)-4-hydroxy-2-((4-(4-methylthiazol-5-yl)benzyl)carbamoyl)pyrrolidin-1-yl)-3,3-dimethyl-1-oxobutan-2-yl)amino)-2-oxoethoxy)ethoxy)acetic acid (CAS: 2172820-08-3)
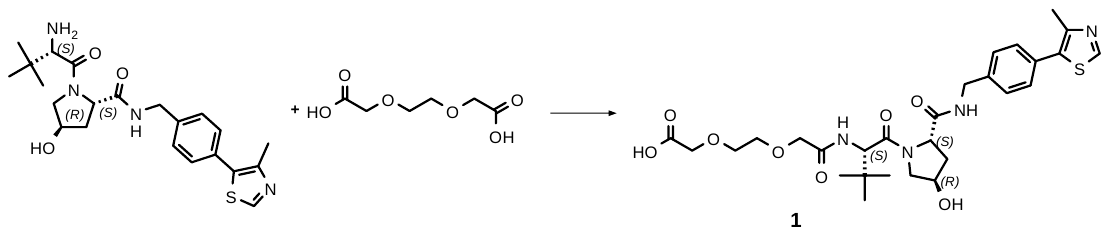

This compound was synthesized according to literature procedure.^[1]^

HPLC-MS: RT = 1.26 min, m/z (M+H)^+^ = 591

Method Info : A: H_2_O + 0.05% HCOOH | B: MeCN + 0.04% HCOOH

T: 45°C | Flow: 3.3 mL/min | MS: 61-1000 amu positive

Column: Chromolith HR RP-18e 50-4.6 mm

1% -> 99% B: 0 -> 2.0 min | 99% B: 2.0 -> 2.5 min

##### Synthesis of Hapten 2 (VHL-6)

(S)-16-((2S,4R)-4-hydroxy-2-((4-(4-methylthiazol-5-yl)benzyl)carbamoyl)pyrrolidine-1-carbonyl)-17,17-dimethyl-14-oxo-3,6,9,12-tetraoxa-15-azaoctadecanoic acid (CAS: 2360516-50-1)

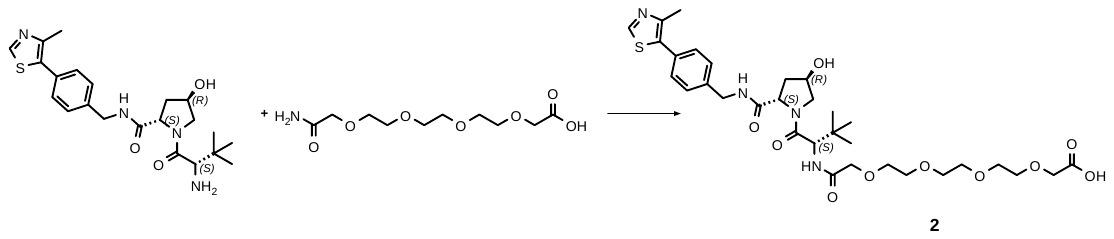

This compound was synthesized according to literature procedure (*MERCK KGAA - WO2023/78813, 2023, A1; Location in patent: Page/Page column 155*).

HPLC-MS: RT = 1.3 min, m/z (M+H)^+^ = 679

Method Info : A: H_2_O + 0.05% HCOOH | B: MeCN + 0.04% HCOOH

T: 40°C | Flow: 3.3 mL/min | MS: 100-2000 amu positive

Column: Chromolith HR RP-18e 50-4.6 mm

0% -> 100% B: 0 -> 2.0 min | 100% B: 2.0 -> 2.5 min

##### Synthesis of Hapten 3 (VHL-7)

(S)-19-((2S,4R)-4-hydroxy-2-((4-(4-methylthiazol-5-yl)benzyl)carbamoyl)pyrrolidine-1-carbonyl)-20,20-dimethyl-17-oxo-3,6,9,12,15-pentaoxa-18-azahenicosanoic acid (CAS: 2172820-13-0)

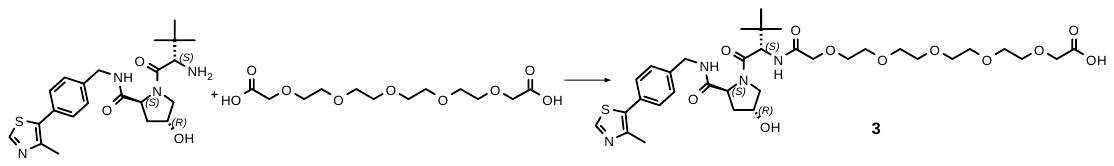

This compound was synthesized according to literature procedure.^[2]^

HPLC-MS: RT = 1.33 min, m/z (M+H)^+^ = 723

Method Info : A: H_2_O + 0.05% HCOOH | B: MeCN + 0.04% HCOOH

T: 40°C | Flow: 3.3 mL/min | MS: 100-2000 amu positive

Column: Chromolith HR RP-18e 50-4.6 mm

0% -> 100% B: 0 -> 2.0 min | 100% B: 2.0 -> 2.5 min

##### Synthesis of 11 (cMETd1)

N1'-(3-fluoro-4-((7-(3-(3-(3-(((S)-1-((2S,4S)-4-hydroxy-2-((4-(4-methylthiazol-5-yl)benzyl)carbamoyl)pyrrolidin-1-yl)-3,3-dimethyl-1-oxobutan-2-yl)amino)-3-oxopropoxy)propoxy)propoxy)-6-methoxyquinolin-4-yl)oxy)phenyl)-N1-(4-fluorophenyl)cyclopropane-1,1-dicarboxamide (CAS: 2230821-69-7)

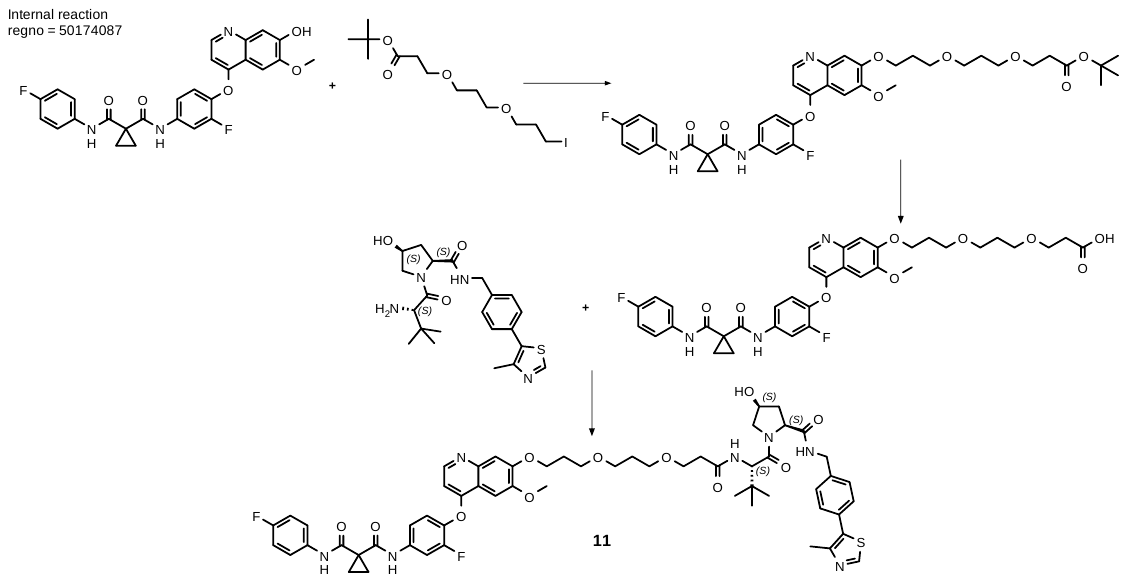

Compound **11** was synthesized according to literature procedure (*ARVINAS - WO2018/226542, 2018, A1, Location in patent: Paragraph 00373*) starting from N-[3-Fluoro-4-[(7-hydroxy-6-methoxy-4-quinolinyl)oxy]phenyl]-N′-(4-fluorophenyl)-1,1-cyclopropanedicarboxamide (CAS: 849217-50-1)

UPLC-MS: HPLC-MS: RT = 0.701 min, m/z (M+2H)^2+^ = 554

System: Waters Acquity UPLC

Method: A: H_2_O + 0,05% HCOOH | B: MeCN + 0.04% HCOOH + 1% H_2_O

T: 40°C | Flow: 0.9 mL/min | Column: Kinetex EVO-C18 1.7 μm 50-2.1 mm

1% -> 99% B: 0 -> 1.0 min | 99% B: 1.0 -> 1.3 min

##### Synthesis of pH responsive dye 15 (VH032-pHAb dye)

1-(9-{2-carboxylato-5-[(2-{2-[2-({[(2S)-1-[(2S,4R)-4-hydroxy-2-({[4-(4-methyl-1,3-thiazol-5-yl)phenyl]methyl}carbamoyl)pyrrolidin-1-yl]-3,3-dimethyl-1-oxobutan-2-yl]carbamoyl}methoxy)ethoxy]ethoxy}ethyl)carbamoyl]phenyl}-6-[4-(4-sulfobutyl)piperazin-1-yl]-3H-xanthen-3-ylidene)-4-(4-sulfobutyl)-1lambda5-piperazin-1-ylium

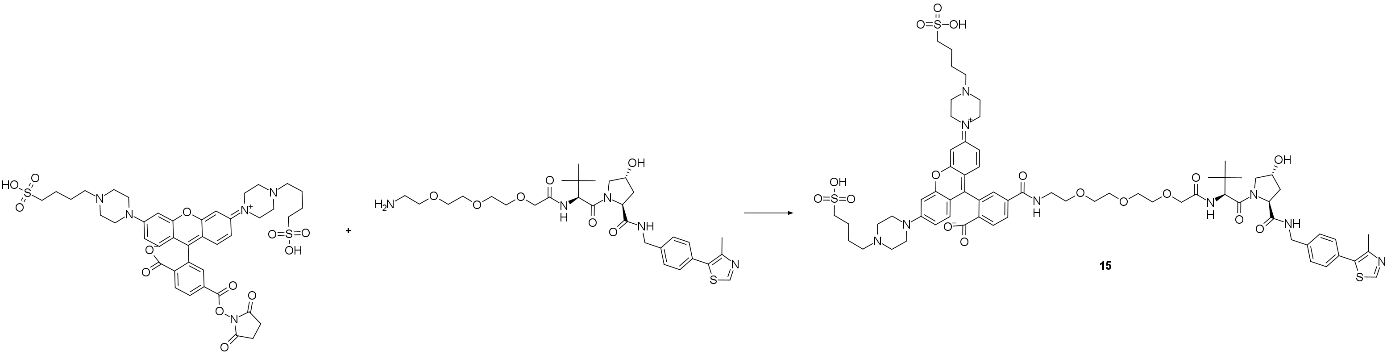

A 0.5 mL septum-sealed microwave vial with stir flea was heated with a heat gun under vacuum and cooled down under a flow of argon gas (needle technique). 1-[9-(2-carboxylato-5-{[(2,5-dioxopyrrolidin-1-yl)oxy]carbonyl}phenyl)-6-[4-(4-sulfobutyl)piperazin-1-yl]-3H-xanthen-3-ylidene]-4-(4-sulfobutyl)-1lambda5-piperazin-1-ylium (3 mg; 3.4 µmol; 1 eq.) (12 x 250 µg dry aliquots) were dissolved in a total of 250 µL dry DMSO. The pink solution was immediately transferred to the reaction vessel, and 1-[9-(2-carboxylato-5-{[(2,5-dioxopyrrolidin-1-yl)oxy]carbonyl}phenyl)-6-[4-(4-sulfobutyl)piperazin-1-yl]-3H-xanthen-3-ylidene]-4-(4-sulfobutyl)-1lambda5-piperazin-1-ylium (3 mg; 3.4 µmol; 1 eq.) was added, followed by *N*-ethyldiisopropylamine (*^i^*Pr_2_NEt) (12 µL; 68 µmol; 20 eq.) as a base. The vessel was sealed and fitted with an argon balloon *via* the septum. The reaction mixture was shielded from light by cardboard boxes and stirred at RT, which led to a deep purple solution. The reaction mixture was stirred overnight at RT, diluted with 750 µL DMSO and purified by preparative RP-HPLC-MS (Sunfire column). Target fractions were lyophilized under light exclusion, giving a pink-purple solid as the product.

HPLC-MS: RT = 1.76 min, m/z (M+2H)^2+^ = 693

Method Info : A: H_2_O + 0.05% HCOOH | B: MeCN + 0.04% HCOOH + 1% H_2_O

T: 40°C | Flow: 1.4 mL/min | MS: 61-1000 amu positive

Column: SunFire C18 5.0 μm 100-3 mm

1% -> 99% B: 0 -> 2.0 min | 99% B: 2.0 -> 2.7 min

##### Synthesis of BETTY2 (9, here: compound 1) and BETTY3 (10, here: compound 2) by ChemPartner, China.

**Synthesis Scheme**

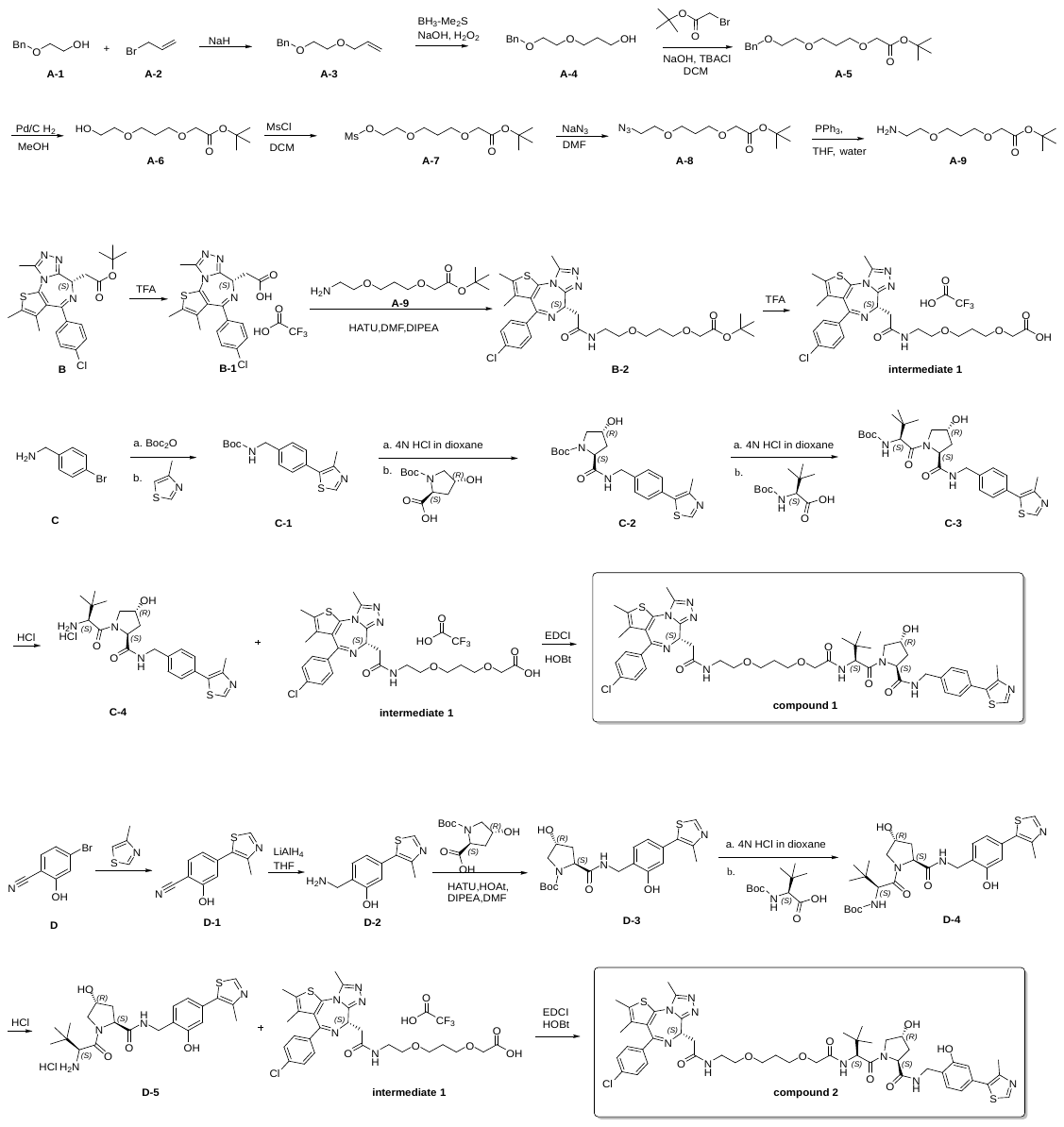

Synthesis of compound A-3

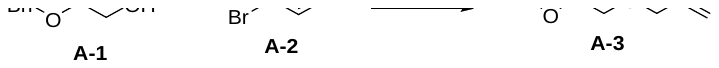

To a solution of compound A-1 (30 g, 197 mmol) in tetrahydrofuran (THF) (400 mL) was carefully added sodium hydride (9.46 g, 237 mmol) and compound A-2 (28.6 g, 237 mmol) at 0 °C. The reaction mixture was stirred at 20 °C for 16 hours, quenched with water (~ 100 mL), partitioned with ethyl acetate (700 mL) and water (200 mL). The organic layer was washed with saturated aqueous sodium chloride (100 mL), dried with anhydrous sodium sulfate, filtered and evaporated to give compound A-3 as yellow oil (35 g, yield: 91%).

Synthesis of compound A-4

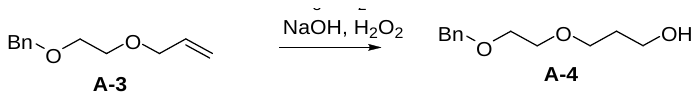

To a mixture of compound A-3 (35 g, 182.3 12.09 mmol) in THF (200 mL) was added borane dimethylsulfide (BH_3_-Me_2_S) (10 M, 27 mL) dropwise at 0 °C under nitrogen (N_2)_ and the resulting mixture was stirred at RT for 2 hours. TLC indicated starting material was consumed completely and one major new spot was detected. Then H_2_O (124 mL), NaOH (3 M, 14 mL) and H_2_O_2_ (146 g, 1292.5 mmol, 124 mL, 30% wt) were added sequentially to this reaction at 0 °C and the resulting mixture was allowed to stir another 2 hours at 20 °C. TLC indicated starting material was consumed completely. The reaction mixture was quenched with saturated Na_2_CO_3_ (150 mL) and extracted with EtOAc (150 mL x 3). The organic layer was dried over Na_2_SO_4_, filtered and concentrated. The residue was purified by column chromatography (SiO_2_, Petroleum ether/Ethyl acetate=10/1 to 1/1) to afford compound A-4 as a colorless oil (16 g, yield: 41%).

Synthesis of compound A-5

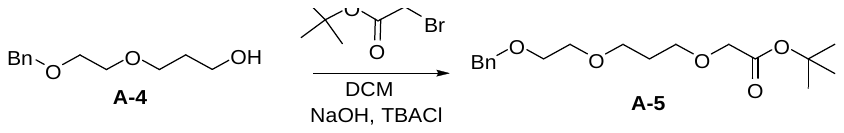

A mixture of compound A-4 (16 g, 76 mmol), tert-butyl bromoacetate (45.0 mL, 304 mmol), tetra-n-butylammonium chloride (21.15 g, 76 mmol) and NaOH (76 mL, 761 mmol) in DCM (200 mL) was stirred vigorously for 18 hours. The mixture was concentrated and partitioned between 1000 mL each of ethyl acetate and water. The organic layer was washed with 200 mL of saturated aqueous sodium chloride, dried over anhydrous sodium sulfate, filtered and concentrated. The residue, combined with above batch, was purified by chromatography (silica, 5-100% ethyl acetate in petroleum ether afforded compound A-5 as light yellow oil (14 g, yield:55%).

Synthesis of compound A-6

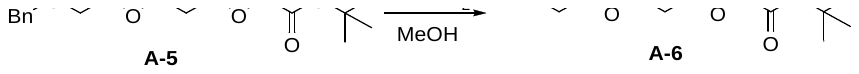

To the mixture of compound A-5 (14 g, 43.2 mmol) in MeOH (150 ml) was added Pd-C (2.76 g, 2.59 mmol) under nitrogen and then the reaction was degassed with hydrogen and stirred under hydrogen for 16 hours. LCMS showed that all the SM had disappeared. The reaction, combined with above batch, was filtered through celite and washed with ethyl acetate. The solvent was removed to give compound A-6 which was used into next step directly (10 g, yield: 94%).

Synthesis of compound A-7

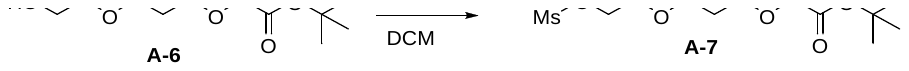

To a solution of compound A-6 (10 g, 42.7 mmol) and TEA (triethylamine) (11.90 mL, 85 mmol) in DCM (300 mL) was added Ms-Cl (methanesulfonyl chloride) (4.99 mL, 64.0 mmol) slowly at 0 °C. Then the reaction mixture was stirred for 3 hr. TLC showed that the starting material had disappeared and a new spot was observed. The reaction mixture was quenched by addition of water (~ 100 mL), then extracted with dichloromethane (100 mL). The combined organic phase was washed with brine (50 mL), dried over sodium sulfate, filtered and concentrated under reduced pressure to afford compound A-7 as colorless liquid, which was used directly for the next step (13 g, yield: 93%).

Synthesis of compound A-8

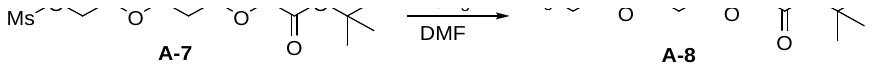

To a solution of compound A-7 (13 g, 41.6 mmol) in DMF (90 ml) was added sodium azide (3.25 g, 49.9 mmol). Then the reaction mixture was stirred at 70 °C for 16 hours. Then water (300 mL) was added and the reaction mixture was extracted with ethyl acetate (200 mL x 2). The combined organic phases were washed with brine (100 mL), dried over sodium sulfate, filtered and concentrated under reduced pressure to give a residue, which was purified by column chromatography eluted with petroleum ether : ethyl acetate = 100:1 to 3:1 to afford compound A-8 as colorless oil (10 g, yield: 93%).

Synthesis of compound A-9

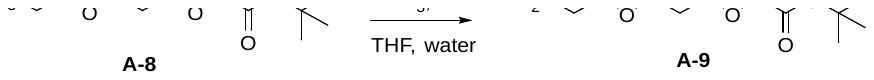

A mixture of compound A-8 (10 g, 36.6 mmol) and triphenylphosphine (14.41 g, 55.0 mmol) in THF (150 ml) / Water (4.55 ml) was stirred at 20 °C for 16 hours. Thin-layer chromatography (TLC) (ninhydrin, 5% methanol in dichloromethane) indicated the reaction was complete. The solvent was removed and the residue was loaded onto silica gel and purified by chromatography (silica, 1-10% methanol/ammonia in dichloromethane) to give compound A-9 as light yellow oil (7.5 g, yield:83%).

Synthesis of compound B-1

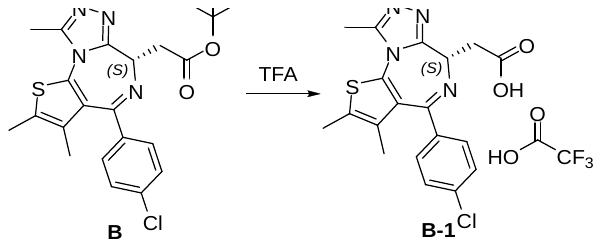

Compound B (5 g, 10.75 mmol) was dissolved in TFA (20 mL). The mixture was stirred at 60 °C for 1 hour. The mixture was concentrated under reduce pressure to afford crude compound B-1 as an yellow solid, which was used in the next step without further purification (6.6 g, TFA salt).

**Synthesis of compound B-2**

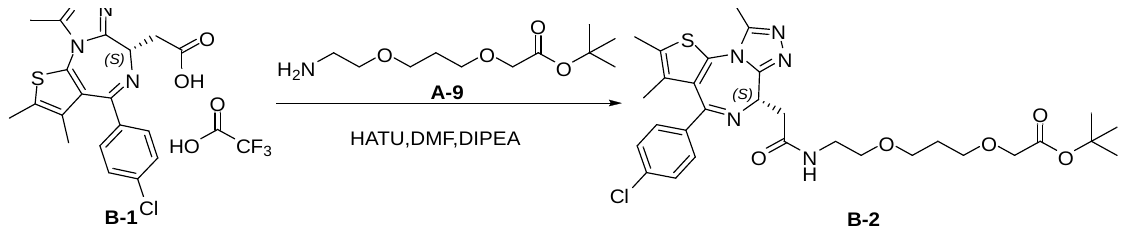

To a solution of compound B-1 (1.2 g, 3.0 mmol), compound A-9 (1.05 g, 4.5 mmol) and HATU (1.7 g, 4.5 mmol) in DMF (15 mL) was added DIPEA (1.56 mL, 9.0 mmol). The mixture was stirred at RT for 1 hour then purified by Reversed-phase chromatography (C18) (0.1% NH_4_HCO_3_ in water, 0-95%MeCN) to give compound B-2 as a white solid (1.2 g, yield: 67%).

**Synthesis of intermediate 1**

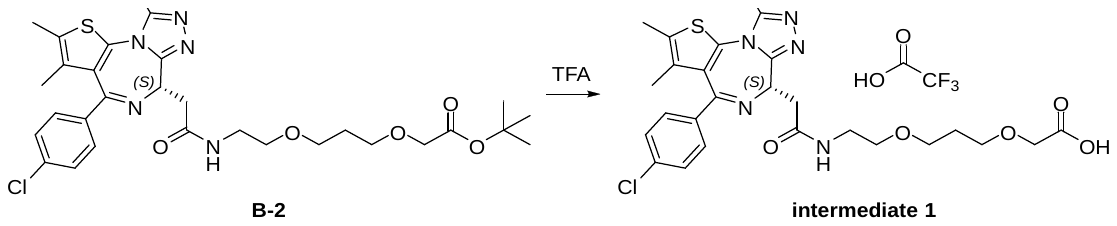

To a solution of compound B-1 (1.2 g, 2.0 mmol) in DCM (10 mL) was added TFA (5 mL) and the mixture was stirred at RT for 2 hours, then the reaction mixture was concentrated under reduced pressure and purified by Reversed-phase chromatography (C18) (0.1% TFA in water, 0-95%MeCN) to give compound intermediate 1 as a yellow solid (950 mg, yield: 73%).

Synthesis of compound C-1

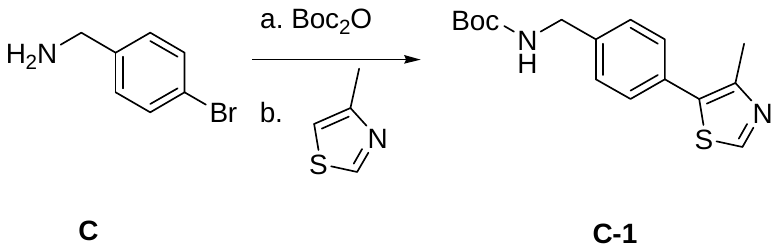

To a mixture of compound C (11.2 g, 60.3 mmol) and NaHCO_3_ (4.0 g, 48.2 mmol) in water (10 mL) and ethyl acetate (60 mL), (Boc)_2_O (15.7 g, 72.4 mmol) at 5 °C was added. The reaction was stirred for 2 hours, TLC showed the reaction was complete. The reaction mixture was filtered. The solid fraction was collected and suspended in a mixture of hexane (40 mL) and water (10 mL) for 0.5 h. The mixture was filtered, and the solid fraction was collected and evaporated to dryness at 50 °C to afford the title compound as a white solid. The solid was dissolved in DMF (40 mL), then 4-methylthiazole (11.9 g, 120.6 mmol), palladium (II) acetate (137 mg, 0.60 mmol), and potassium acetate (11.8 g, 120.6 mmol) was added. The mixture was stirred at 90 °C under nitrogen for 18 h. After cooling to ambient temperature, the reaction mixture was filtered. 50 mL of water was added to the filtrate, and the resulting mixture was stirred at ambient temperature for 4 h. The reaction mixture was filtered. The solid was collected by filtration and dried in an oven at 50 °C to afford compound C-1 as a gray solid (15.3 g, yield: 83%).

Synthesis of compound C-2

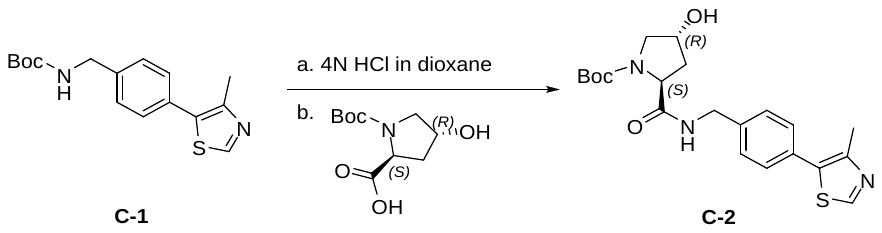

Compound C-1 (15.3 g, 50.3 mmol) was dissolved in 4 N HCl in dioxane (38 mL, 150.9 mmol) and MeOH (35 mL), and the mixture was stirred at ambient temperature for 12 h. The mixture was concentrated, and the residue was dried under vacuum to afford the intermediate. HATU (22.9 g, 60.3 mmol) was added to a solution of this intermediate, (2S,4R)-1-(tert-Butoxycarbonyl)-4-hydroxypyrrolidine-2-carboxylic acid (11.6 g, 50.3 mmol), and DIPEA (26 mL, 150.9 mmol) in DMF (100 mL) at 0 °C under N_2_. The mixture was stirred at ambient temperature for 12 hours. TLC showed that the reaction was complete. The reaction mixture was quenched with H_2_O (200 mL) and extracted with EtOAc (150 mL x 2). The combined organic layer was washed with brine (200 mL) and dried over Na_2_SO_4_. The organic solution was filtered and concentrated. The residue was purified by silica gel flash column chromatography with hexane:EtOAc (100:1−1:100), then DCM:MeOH (10:1) to afford the compound C-2 as white solid (18.1 g, yield: 86%).

Synthesis of compound C-2

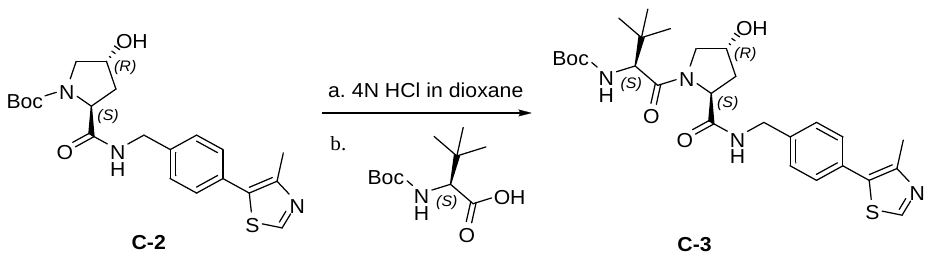

Compound C-2 (16.0 g, 38.3 mmol) was dissolved in 4 N HCl in dioxane (38 mL, 153.2 mmol) and MeOH (38 mL), and the mixture was stirred at ambient temperature for 12 hours. The mixture was then concentrated, and the residue was dried under vacuum to afford intermediate, HATU (21.8 g, 57.4 mmol) was added to a solution of this intermediate, (S)-2-((tert-butoxycarbonyl)amino)-3,3-dimethylbutanoic acid (8.8 g, 38.3 mmol), and DIPEA (20 mL, 114.9 mmol) in DMF (100 mL) at 0 °C under N_2_. The mixture was stirred at ambient temperature for 12 hours when TLC showed that the reaction was complete. The reaction mixture was quenched with H_2_O (200 mL) and extracted with EtOAc (150 mL x 2). The combined organic layer was washed with brine (100 mL) and dried over Na_2_SO_4_. The organic solution was filtered and concentrated. The residue was purified by silica gel flash column chromatography with DCM:MeOH (10:1) to afford the desired compound C-3 as an off-white solid (16.0 g, yield: 79%).

Synthesis of compound C-4

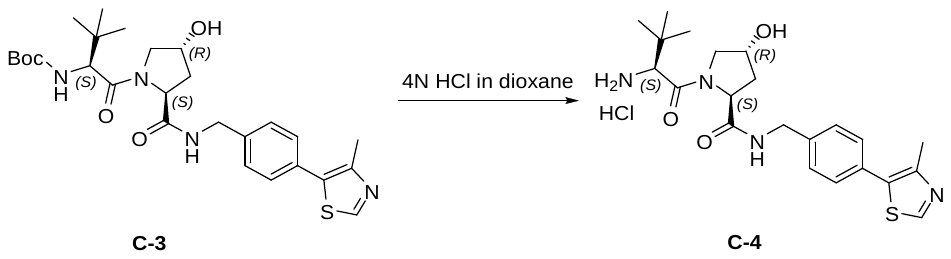

Compound C-3 (4.0 g, 7.55 mmol) was dissolved in 4 N HCl in dioxane (9.0 mL) and MeOH (9.0 mL), and the mixture was stirred at ambient temperature for 12 h. The mixture was then concentrated, and the residue was dried under vacuum to afford crude compound C-4 as an off-white solid, which was used in the next step without further purification. (4.5 g, HCl salt).

**Synthesis of compound 1**

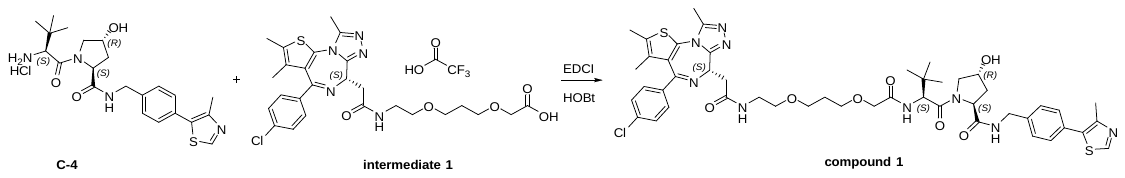

To a solution of compound C-4 (50 mg, HCl salt), intermediate 1 (50 mg, 0.09 mmol), EDCI (27 mg, 0.14 mmol) and HOBt (19 mg, 0.14 mmol) in DMF (1.0 mL) was added DIPEA (0.05 mL, 0.27 mmol). The mixture was stirred at RT for 1 hour then purified by Reversed-phase chromatography (C18) (0.1% NH_4_HCO_3_ in water, 0-95% MeCN) to give compound 1 as a white solid (35 mg, yield: 43%).

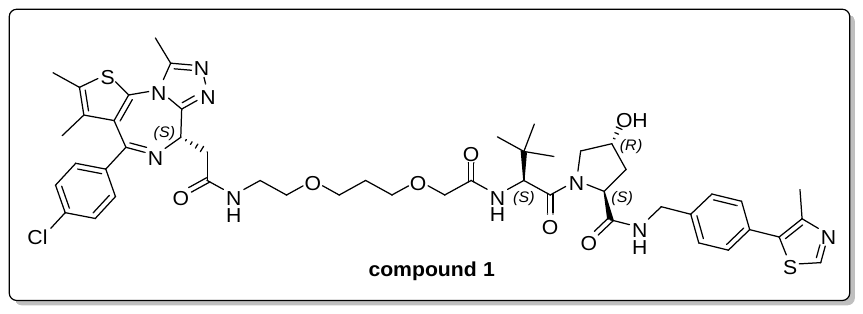

LC/MS (ESI): m/z = 486.7 [M+H]^+^ 1/2. RT = 1.74 min

^1^H NMR (400 MHz, MeOD-*d*4) δ 8.88 (d, *J* = 6.5 Hz, 1H), 7.43 (ddd, *J* = 11.1, 9.8, 5.4 Hz, 8H), 4.74 – 4.33 (m, 6H), 4.07 – 3.78 (m, 4H), 3.75 – 3.54 (m, 6H), 3.45 (dt, *J* = 12.5, 7.4 Hz, 3H), 2.70 (s, 3H), 2.47 (t, *J* = 5.6 Hz, 6H), 2.30 – 1.85 (m, 4H), 1.71 (s, 3H), 1.04 (d, *J* = 8.8 Hz, 9H).

**Synthesis of compound D-1**

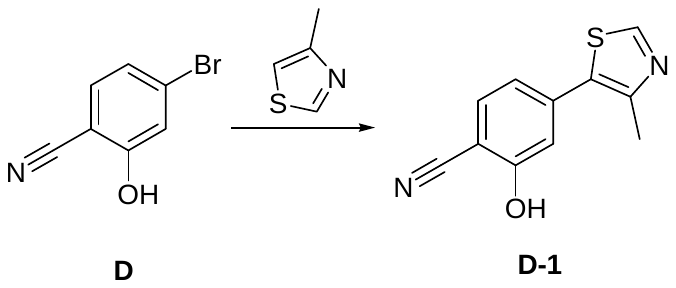

A mixture of compound D (15.0 g, 76.1 mmol) in DMF (40 mL), then 4-methylthiazole (15.1 g, 152.2 mmol), palladium (II) acetate (853 mg, 3.80 mmol), and potassium acetate (14.9 g, 152.2 mmol) was added. The mixture was stirred at 90 °C under nitrogen for 18 hours. After cooling to ambient temperature, the reaction mixture was filtered. Fifty milliliters of water was added to the filtrate, and the resulting mixture was stirred at ambient temperature for 4 hours. The reaction mixture was filtered. The solid was collected by filtration and dried in an oven at 50 °C to afford compound D-1 as a yellow solid (12.3 g, yield: 75%).

**Synthesis of compound D-2**

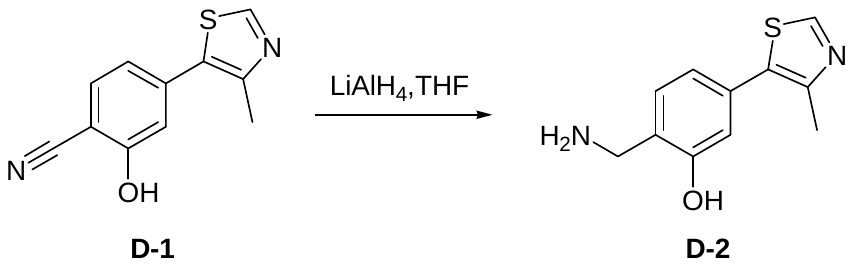

To a solution of compound D-1 (12.3 g, 56.9 mmol ) in THF (60 mL) was added LiAlH_4_ (1 M in THF, 142 mL, 142 mmol) at 0 °C. The reaction mixture was stirred at 50 °C for 30 min, and then THF (100 mL) and 15% NaOH (5.6 mL) was added at 0 °C. Na_2_SO_4_ was added to the mixture and the mixture was stirred at room temperature for 15 min. The slurry was filtered over a short pad of Celite and rinsed with THF (3 x 50 mL), and 20% MeOH in CH_2_Cl_2_ (3 x 50 mL), then the filtrate was concentrated under reduced pressure. The residue was purified by column chromatography on silica gel (0-10% MeOH/NH_3_ in CH_2_Cl_2_) to give compound D-2 as a colorless oil (6.2 g, yield: 50%).

Synthesis of compound D-3

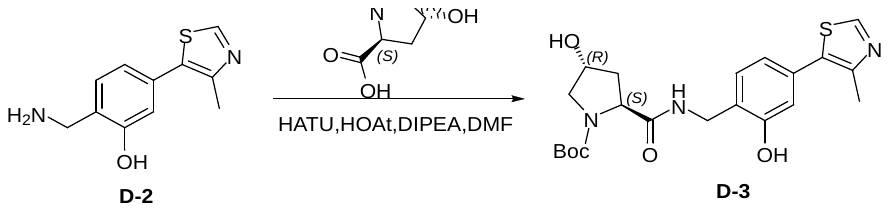

To a solution of compound D-2 (6.2 g, 28.2 mmol), (2S,4R)-1-(tert-Butoxycarbonyl)-4-hydroxypyrrolidine-2-carboxylic acid (6.5 g, 28.2 mmol), HATU (10.7 g, 28.2 mmol) and HOAt (3.8 g, 28.2 mmol) in DMF (20 mL) was added DIPEA (9.8 mL, 56.4 mmol) at 0 °C under N_2_. The mixture was stirred at ambient temperature for 12 hours. TLC showed that the reaction was complete. The reaction mixture was quenched with H_2_O (200 mL) and extracted with EtOAc (150 mL x 2). The combined organic layer was washed with brine (200 mL) and dried over Na_2_SO_4_. The organic solution was filtered and concentrated. The residue was purified by column chromatography on silica gel (1-10 % MeOH in CH_2_Cl_2_) to afford the compound D-3 as white solid (8 g, yield: 65%).

Synthesis of compound D-4

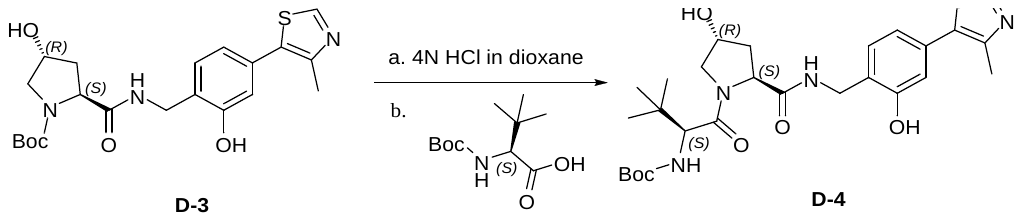

Compound D-3 (8.0 g, 18.5 mmol) was dissolved in 4 N HCl in dioxane (18 mL) and MeOH (18 mL), and the mixture was stirred at ambient temperature for 12 hours. The mixture was then concentrated, and the residue was dried under vacuum to afford the intermediate, HATU (10.5 g, 27.7 mmol) was added to a solution of this intermediate, (S)-2-((tert-butoxycarbonyl)amino)-3,3-dimethylbutanoic acid (4.3 g, 18.5 mmol), and DIPEA (9.6 mL, 55.5 mmol) in DMF (50 mL) at 0 °C under N_2_. The mixture was stirred at ambient temperature for 12 hours when TLC showed that the reaction was complete. The reaction mixture was quenched with H_2_O (200 mL) and extracted with EtOAc (150 mL x 2). The combined organic layer was washed with brine (100 mL) and dried over Na_2_SO_4_. The organic solution was filtered and concentrated. The residue was purified by silica gel flash column chromatography with DCM:MeOH (10:1) to afford the desired compound D-4 as an off-white solid (5 g, yield: 50%).

Synthesis of compound D-5

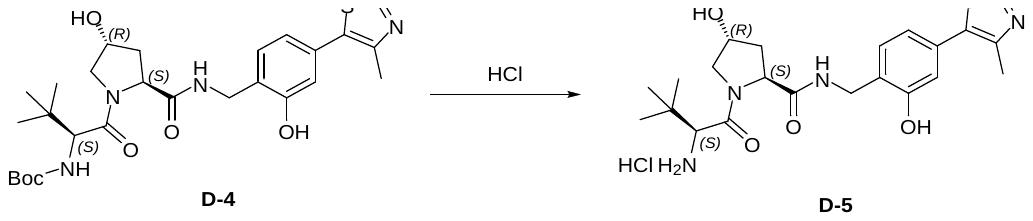

Compound D-4 (3.5 g, 6.4 mmol) was dissolved in 4 N HCl in dioxane (9.0 mL) and MeOH (9.0 mL), and the mixture was stirred at ambient temperature for 12 hours. The mixture was then concentrated and the residue was dried under vacuum to afford crude compound D-5 as an off-white solid, which was used in next step without further purification (4.0 g, HCl salt).

**Synthesis of compound 1**

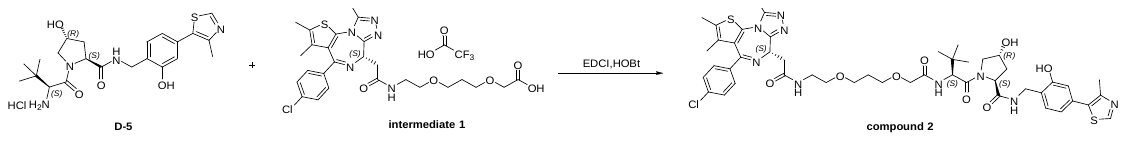

To a solution of compound D-5 (301 mg, HCl salt), intermediate 1 (350 mg, 0.52 mmol), EDCI (149 mg, 0.78 mmol) and HOBt (105 mg, 0.78 mmol) in DMF (8 mL) was added DIPEA (0.27 mL, 1.56 mmol). The mixture was stirred at RT for 1 hour, then purified by Reversed-phase chromatography (C18) (0.1% NH_4_HCO_3_ in water, 0-95% MeCN) to give compound 1 as a white solid (200 mg, yield: 39%).

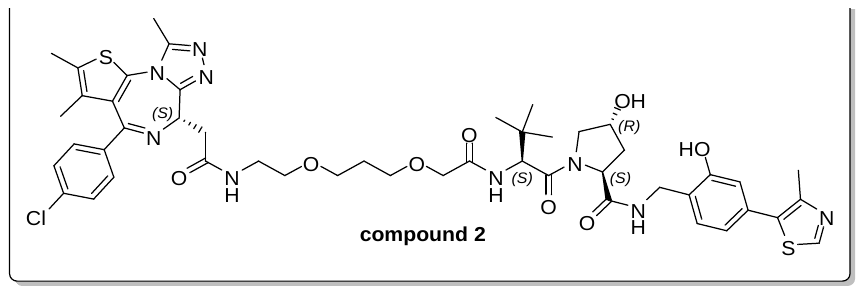

LC/MS (ESI): m/z = 494.7 [M+H]^+^ 1/2. RT = 1.77 min

^1^H NMR (400 MHz, MeOD-*d*4) δ 8.86 (d, *J* = 6.1 Hz, 1H), 7.62 – 7.20 (m, 5H), 7.00 – 6.82 (m, 2H), 4.81 – 4.25 (m, 6H), 3.99 (q, *J* = 15.4 Hz, 2H), 3.83 (dt, *J* = 11.1, 7.4 Hz, 2H), 3.70 – 3.55 (m, 6H), 3.53 – 3.40 (m, 3H), 2.71 (d, *J* = 4.6 Hz, 3H), 2.48 (t, *J* = 7.0 Hz, 6H), 2.27 – 2.06 (m, 2H), 1.91 (p, *J* = 6.2 Hz, 2H), 1.72 (d, *J* = 5.9 Hz, 3H), 1.10 – 0.95 (m, 9H).

##### Conjugation of VH032-based haptens

For the preparation of immunogens and screening compounds, VH032-based haptens (**1** (VHL-1), **2** (VHL-6), **3** (VHL-7), **4** (VHL-c)) were dissolved separately in conjugation buffer (0.1 M MES, 0.9 M NaCl, 0.02% sodium azide; pH 4.7) to a final concentration of 4 mg/mL and mixed either with a solution of 10 mg/mL keyhole limpet hemocyanin (KLH), 10 mg/mL cationic Bovine Serum Albumin (cBSA) or 7 mg/mL human Fc (fragment crystallizable) at a final protein/hapten molar ratio of 1:100. To this mixture, a 10 mg/mL aqueous solution of 1-ethyl-3-(3-dimethyllaminopropyl) carbodiimide (EDCI) was added (final protein-to-EDCI molar ratio 1:1750) and the reaction was incubated overnight. Reaction mixes were purified using Zeba Spin Desalting Columns that were pre-equilibrated in phosphate buffered saline (PBS, 0.137 M NaCl, 0.0027 M KCl, 0.01 M Na_2_HPO_4_, 0.0018 M KH_2_PO_4_, pH 7.4) (7K MWCO, Thermo Scientific). Protein concentrations were determined with Bradford reagent using unconjugated cBSA, KLH or human Fc as standards. The hapten/protein ratio for cBSA and human Fc conjugates was analyzed by MALDI-MS.

#### Methods

##### Immunization and cDNA preparation

Livestock farming and immunization of New World Camelids (llamas, alpacas, huarizos) was managed by Preclinics GmbH (Potsdam, Germany) according to national and international guidelines. The study was approved by an Institutional Animal Care and Use Committee, the Lower Saxony State Office for consumer protection and food safety (Oldenburg, Germany, reference number 33.19-42502-05-17A210). For generating an immune library, the three llamas (Lama glama) „Emma“, „Ferdinand“, and „Elvis“ were immunized subcutaneously with 300 µg mixed 1:1:1 VHL-1/-6/-7-carrier conjugate (100 µg of each VHL-carrier conjugate) in 0.5 mL PBS mixed with an equal volume of GERBU adjuvant S (GERBU Biotechnik GmbH, Heidelberg, Germany). Four weeks after priming, all animals received five boost immunizations identical to the prime injection in a two-week interval. For the immunization, KLH and cBSA were used alternatingly as carrier proteins for the VHL haptens, i. e. for injection 1, and 3 KLH conjugates and for injections 2, and 4 to 6 cBSA conjugates were used. In order to monitor the development of immune responses, serum samples were gained directly prior to prime immunization and on day 35, 49, and 77 of the immunization project. Four days after the sixth injection (day 88), a final blood sample was gained by venipuncture together with a final serum sample. Subsequently, peripheral blood mononuclear cells (PBMCs) were isolated by density gradient centrifugation on lymphocyte separation medium (LSM, Corning Inc., Corning, NY, USA) with a density of 1.077 to 1.080 g/mL at 20°C. The isolated cells from the interphase were collected and washed in DPBS (Corning). Finally, the cells were counted and lysed in RA1 buffer (Macherey-Nagel, Düren, Germany), 15 mM DTT and stored frozen until subsequent RNA and cDNA preparation. Humoral immune responses were monitored in diluted serum or plasma by ELISA on 96 well microtiter plates (high binding half area, Corning Inc., Corning, NY, USA) coated with VHL-1, VHL-7, or VHL-c human Fc conjugates and human IgG, respectively, as control. All proteins were diluted in 50 mM carbonate buffer pH 9.6 to a final concentration of 5 µg/mL. From each serum sample, a 1:4 serial dilution starting from 1:25 in PBS/BSA (1% w/v) was prepared. For detection of total IgG, dilutions starting from 1:100 were analyzed. Bound total camelid IgG and heavy-chain IgG, respectively, was detected using peroxidase-conjugated monoclonal mouse-anti-llama IgG antibodies (produced in collaboration with University of Potsdam, PMID: 29475500, DOI: 10.1016/j.vetimm.2018.01.006), and 3,3’,5,5’-tetramethylbenzidine (TMB One, Kementec, Taastrup, Denmark) as substrate. The absorbance at 450 nm and 620 nm reference wavelength was measured using a Mithras LB940 multimode reader (Berthold Technologies, Bad Wildbad, Germany). RNA was isolated from the lysed PBMCs using NucleoSpin RNA Midi kit for RNA purification (Macherey-Nagel, Düren, Germany) according to the manufacturer’s instructions. The RNA was eluted in RNase-free water. For preparation of cDNA, RevertAid H Minus First Strand cDNA Synthesis Kit (K1632, Thermo Fisher Scientific Inc.) was used according to the manufacturer’s instructions. For each animal, ten cDNA reactions using both oligo(dT)18 and random hexamer primers were used and pooled afterwards individually.

##### Generation of antibody phage libraries

The selection of VHL specific antibodies from a VHH antibody immune library generated from immunized llamas was managed by YUMAB GmbH (Braunschweig, Germany). Therefore, the cDNA pool was used for the amplification of the VHH gene sequences by PCR using proprietary gene specific primer pairs. In the first PCR, primers were used that bind to the leader peptide and CH2 region of the Llama Ig-heavy chain. Using gel extraction, gene sequences that are lacking the CH1 region were extracted and separated from gene amplicons that contain the CH1 domain. After purification, VHH genes were amplified in a second PCR. Here, the primers bind directly to the beginning and end of the VHH domain. Additionally, these primers contain a SapI restriction site, which allows Golden Gate cloning into YUMAB’s phage display vector. As a result, the VHH will be genetically fused in-frame to a Myc- and HIS-tag and to a truncated pIII gene encoded on the vector backbone. The tagged VHH is separated from the pIII by a Trypsin cleavage site and an Amber stop codon. This allows Trypsin specific elution of antibody-phage during panning by cleaving the VHH part from the phage surface. Additionally, the Amber Stop codon allows soluble VHH production for screening purposes, when an amber suppressor E. coli strain is used. After purification of the amplified VHH gene sequences, cloning was done into YUMAB’s antibody-phage display vector. For that purpose, the vector and VHH amplicons were mixed into a golden gate reaction, including T4 reaction buffer, T4 ligase and SapI. A total of 30 restriction and ligation cycles were performed. After heat inactivation for 20 min at 70°C, the ligation products were desalted using Amicon spin columns (30 K). The desalted DNA was mixed with ER2738 Electrocompetent Cells (Lucigen) and incubated for up to 5 min on ice, before electroporation was performed (1.7 kV pulse). The bacteria were recovered in SOC medium and incubated for 1 h at 37°C at 650 rpm. The size of the antibody-gene library was determined by serial dilution and colony counting of a sample of the recovered bacteria. For that purpose, the bacteria were serially diluted in LB medium and transferred onto 2YT agar plates, containing ampicillin (100 µg/mL) and glucose (0.1 M) (2YT-GA). After overnight incubation at 37°C, the number of colonies was counted visually, and the total number of recovered clones calculated. Additionally, the single clones were used for determination of the VHH insert-rate by cPCR and the number of clones with a functional ORF determined by DNA-sequence analysis. Simultaneously, the remaining recovered bacteria were incubated in 2YT-GA and incubated at 37°C and 250 rpm until an OD600 of 0.5 was reached. A sample of the culture was extracted, and bacteria were infected with M13K07 helper-phage (MOI of 20) for 30 min at 37°C. After another propagation cycle for 30 min at 37°C and 250 rpm, the infected bacteria were pelleted by centrifugation at ~3000 g for 10 min. For the production of antibody-phage particles, the bacteria were resuspended in glucose-free 2YT medium, containing Ampicillin (100 µg/mL) and Kanamycin (50 µg/mL) (2YT-AK). The cultivation was continued for 16 h at 30°C. For the purification of the antibody-phage particles, the culture was centrifuged for 40 min at ~3000 g to separate the bacteria from the antibody-phage containing culture supernatant. Then, PEG/NaCl precipitation of the antibody-phage particles was performed, followed by centrifugation (1 h at ~3000 g) and resuspension of the pelleted antibody-phage in phage-dilution buffer. The phage preparation was polished with a high-speed centrifugation step (2 min at ~16000 g). The supernatant was recovered and used for antibody-phage selection. After purification of the antibody-phage particles, the phage concentration was determined. For that purpose, the phages were serially diluted in LB medium and used for the infection of E. coli (30 min at 37°C). The infected bacteria were transferred onto 2YT agar plates, containing ampicillin (100 µg/mL) and glucose (0.1 M) (2YT-GA). After overnight incubation at 37°C, the number of colonies was counted visually, and the concentration of colony forming units calculated. The size of the antibody phage library added up to a diversity of 1.44 x 10^9^ colony forming units. In addition, the insert-rate was determined by colony PCR and the number of clones with a functional ORF determined by DNA-sequence analysis. Additionally, the presence of an antibody-pIII fusion protein was checked by SDS-PAGE, western blotting and anti-pIII immunoblot staining of the antibody-phage particles.

##### Phage display and library screening

Phage display and screening was conducted at Yumab GmbH, Germany. First, the library was cleared from unspecific or cross-reactive antibody-phages. For that purpose, antibody-phage particles (50-100 x fold excess of the library diversity) were diluted in 2% BSA solution, containing 0.05% Tween20 (2% BSA-PBST) and incubated first in Streptavidin-coated ELISA plates (coated with 10 µg/mL, 200 µL/well) for 1 h at RT, followed by incubation with magnetic Streptavidin beads (Dynabeads, Thermo Scientific, 50 µL) for at least 1 h at RT under rotation. Antibody-phages that bound to the negative antigens were removed from further selection by isolation of the phage-containing supernatant. After that, the cleared library was selected for target antigen specific antibodies. For that purpose, a conjugate of the von Hippel-Lindau (VHL)-recruiting ligand and a PEGylated crosslinker with pendant amine was acquired ((S,R,S)-AHPC-PEG_3_-NH_2_ hydrochloride Sigma-Aldrich, 901511-50MG) and biotinylated via Biotin-NHS-ester coupling. The biotinylated VHL ligand was purified and analyzed by HPLC. First, the biotinylated VHL ligand (50 nM) was added to the cleared library. Antibody-phages that bound to the biotinylated target antigen were captured and recovered from the solution using magnetic streptavidin beads (Dynabeads, Thermo Scientific, 50 µL). The beads were washed three times with a 2% BSA-PBST solution and twice in PBS in order to remove unspecific or weakly bound antibody-phage particles. Antigen-specific antibody phages were eluted from the beads by Trypsin (10 µg/mL) treatment at 37°C for 30 min. The eluted phages were rescued by infection of E. coli (OD600 = 0.5) in 2YT medium for 30 min at 37°C. After propagation for 30 min at 37°C and 650 rpm, infected bacteria were selected by addition of glucose (0.1 M) and ampicillin (10 µg/mL). The cultivation was continued for 30 min at 650 rpm and 37°C before M13K07 helper phage was added (MOI 1:20). The co-infection with helper phage was performed for 30 min at 37°C, followed by incubation for 30 min at 37°C and 650 rpm. For the amplification of antibody-phage particles, the bacteria were pelleted by centrifugation (~3000 g, 10 min) and the culture medium exchanged to glucose-free 2YT-AK before the cultivation was continued for 16 h at 30°C and 650 rpm. The amplified phages were used for two more selection cycles as described above. To identify the best performing antibody variants targeting VH032 (**5**). The selection outputs of rounds three and four were subjected to single clone analyses. After the antibody-phage selection, the binding characteristics of the monoclonal antibody clones were analyzed. 384 single clones were selected from the selection output after panning round two and three and used for VHH antibody expression in bacteria. For that purpose, eluted phages were used for infection of E. coli (as described above) and cultivated on 2YT-GA agar plates. The plates were incubated at 37°C until single colonies were observed. The single colonies were picked and transferred into microtiter plates containing 2YT-GA culture medium and cultivated at 37°C and 250 rpm for 16 h. For VHH production, 15 µL of each overnight culture was transferred to microtiter plates containing 2YT-A and 50 µM IPTG. Antibody production was done for 16 h at 30°C and 250 rpm. Each antibody production was tested for their binding specificity by ELISA on Streptavidin + biotinylated VHL ligand, Streptavidin, Human Fc-VHL-1, and Human Fc. For that purpose, 384-well ELISA plates were either coated with Streptavidin (40 ng/well), followed by blocking with 2% BSA-PBST solution and capturing of the biotinylated antigen (20 ng/well), or by direct immobilization of the non-biotinylated antigen (20 ng/well), followed by blocking with 2% BSA-PBST. The antibody-containing E. coli productions were mixed with 2% BSA-PBST in a 1:1 ratio, transferred to the washed ELISA plates (30 µL/well) and incubated for 1 h at RT. After washing, bound antibodies were detected via a secondary anti-Myc HRP conjugated antibody. After washing, binding was quantified by standard TMB staining and absorbance reading. Antibody clones were identified as antigen specific, if the ELISA binding signal to the positive antigens was ≥ 0.1, the ELISA binding signal to the negative antigens was ≤ 0.1, and the S/N ratio between positive and negative antigens was ≥ 10.

562 Hits were identified from the llama library. All Hits were used for DNA-sequence analysis to identify antibodies with a unique antibody sequence (≥ 1 amino acid difference in the CDRs). 113 unique clones were identified from the llama library. To identify clones with the best binding affinity, a biolayer interferometry (BLI) off-rate measurement was performed. First, VHH containing culture supernatants were produced of the unique clones. After that, the association and dissociation of the antibody fragment to the biotinylated VHL was measured which was immobilized onto BLI Streptavidin sensors. The binding curve was fitted with a 1:1 binding model and the dissociation rate calculated. Based on the off-rates and antibody sequence information, 10 lead clones (named MIC5 – MIC14) were selected for conversion into the final antibody format.

##### Reformatting of VH032-binding VHH domains into PROxAb Shuttles

Reformatting was conducted at Yumab GmbH, Germany. For this, the VHH genes were amplified from the phagemid DNA by PCR and cloned into two different IgG expression vectors. These expression vectors encoded the therapeutic anti-CD33 antibody gemtuzumab bearing a receptor-silenced human IgG4 PG-SPLE or human IgG1 PG-LALA (only for Shuttles with humanized VHHs) backbone (αCD33)^[3]^, the epidermal growth factor receptor-targeting (EGFR) antibody cetuximab bearing a receptor-silenced human IgG1 PG-LALA backbone (αEGFR) or an antibody targeting digoxigenin bearing either the above-mentioned receptor silenced human IgG4 or IgG1 backbone (αDIG). For vector generation, the individual VHH antibody fragments were genetically fused to the C-terminus of the IgG antibody heavy chains. The VHH domain was separated from the IgG heavy chain via a short glycine-serine amino acid linker (GSGGGSGGSGGGGSG). After sequence verification and preparation of transfection-grade DNA, human embryonic kidney (HEK) cells were transiently transfected with the expression vectors and incubated for 7 days at 37°C and 5% CO_2_ to enable production of the final PROxAb Shuttle antibodies. The cells were separated from the antibody supernatant by centrifugation at 3000 rpm for 20 minutes at 4°C and purified using protein A affinity chromatography. After clearance of the culture supernatant from cells by centrifugation, the IgG antibodies were purified by Protein A affinity chromatography. After adjustment of the buffer to PBS, the protein concentration of the antibodies was determined by UV/VIS spectrometry. Integrity and purity of the antibodies were assessed by SDS-PAGE under reducing conditions. The functional binding activity of the antibodies to the target antigen was measured by ELISA.

***In vitro* characterization of VH032-binding VHH antibody domains**

The kinetic and affinity parameters of the VHH/PROTAC interactions were evaluated by surface plasmon resonance (SPR). αCD33xMIC7, αEGFRxMIC7 or αCD33xhMIC7_1.6 were immobilized onto a high-capacity amine sensor chip (Bruker Daltonics) via the standard amine coupling procedure, at 25 °C. Prior to immobilization, the carboxymethylated surface of the chip was activated with 200 mM 1-ethyl-3-(3-dimethylaminopropyl)-carbodiimide and 25 mM N-hydroxy succinimide (NHS) for 10 min. The PROxAb variants were diluted to 10 µg/mL in 10 mM acetate at pH 4.5 and immobilized on the activated surface chip for 7 min, in order to reach 3,000 to 9,000 response units (RU). The remaining activated carboxymethylated groups were blocked with 1 M ethanolamine pH 8 in a 7 min injection step. HBS-N, which consists of 10 mM HEPES pH 7.4 and 150 mM NaCl, was used as the background buffer during immobilization. PROTACs were prediluted in DMSO, diluted 1:50 in running buffer (12 mM phosphate, pH 7.4, 137 mM NaCl, 2.7 mM KCl, 0.05% Tween20, 2% DMSO) and injected at 10 different concentrations using two-fold dilution series, from 1 µM to 0.002 µM. A DMSO solvent correction (1% - 3%) was performed to account for variations in bulk signal and to achieve high-quality data. Interaction analysis cycles consisted of a 300 sec sample injection (30 μL/min; association phase) followed by 900 sec of buffer flow (dissociation phase). All sensorgrams were processed by first subtracting the binding response recorded from the control surface (reference flow-channel), followed by subtraction of the buffer blank injection from the active flow-channel (target protein immobilized). All datasets were fit to a simple 1:1 Langmuir interaction model to determine the kinetic rate constants. The experiments were performed on a SPR-32 PRO (Bruker Daltonics, Bremen, Germany) at 25°C or 37°C and the interactions were evaluated using the provided Sierra Analyser Software (version 3.4.5.).

##### Generation of PROTAC-complexed PROxAb Shuttles

Complexation to create PROTAC-complexed PROxAb Shuttles was performed as follows: 10 µM (final) antibody-VHH fusion were mixed in PBS pH 7.4 with varying equivalents of VHL-based PROTAC in DMSO to achieve the desired loading. PROxAb Shuttles determined for *in vivo* application were complexed at an antibody concentration of 6 mg/mL. For *in vitro* application, Tween-20 was typically added to a final concentration of 0.3% in order to use a Tecan D300e dispenser for PROxAb Shuttle application on cells. The samples were incubated on a ThermoMixer for 3 h at 25°C while shaking at 650 rpm. Note, 3 h incubation was chosen to assure full loading. However, we observed full complexation within minutes (data not shown). No aqueous Tween-20 was added for complex investigation using size-exclusion chromatography, flow cytometric analysis or native SEC-native MS.

##### Isothermal titration calorimetry

ITC measurements were performed with VP-ITC microcalorimeter from MicroCal/Malvern Panalytical (UK). PROxAb Shuttles, His-tagged MIC7-VHH and the respective VHL-ligand compounds were formulated in 30 mM HEPES; 150 mM NaCl; pH 7.4 with optional addition of 0.01% Tween-20 for all titration experiments. The proteins were loaded into the injection syringe with final concentrations of 25 µM for Shuttles and to 50 μM for VHH. VH032 stock solutions of 10 mM in DMSO were diluted to 5 µM concentrations with buffer and loaded into the sample cell. All buffers were adjusted to a final concentration of 1% (v/v) DMSO. Both, the titrate and titrant solutions were degassed prior to loading the calorimeter cell and injection syringe. ITC titrations were conducted at a constant temperature of 303 K. ITC data analysis was performed using the Origin 7-based (OriginLab Cooperation Northampton, USA) calorimetry customization supplied as standard instrument software by MicroCal / Malvern Panalytical (UK). The integrated heat data were fitted with a one-site binding model to determine the apparent values for affinity, enthalpy, and stoichiometry of binding. Binding parameters of reference ligands were monitored, and the binding stoichiometry was employed as reference point for normalizing the concentration of different protein batches.

##### Native size exclusion liquid chromatography – native Mass spectrometry analysis

An eksigent M5 MicroLC system from SCIEX (ON, Canada) connected to a quadrupole time of flight (QTOF) mass spectrometer (MS) (QTOF X500B, SCIEX, ON, Canada) was used for the analysis. The protein samples were separated using a PolyHYDROXYETHYL A capillary (150 x 0.30 mm, 3 µm, 1000 A). The column temperature was set to 25°C and an isocratic gradient with 100 mM ammonium acetate pH 6.8 was run with 10 µL/min for 10 min. A positive time of flight (TOF) MS method was used for analysis. The electrospray voltage of the X500B QTOF instrument was set to 5500 V and the temperature was 400 °C. Full MS scans were acquired over a mass to charge range of 5000 - 7000 with an accumulation time of 1 sec and time bins to sum 120. The ion source 1 and 2 gases were set to 60 psi. The declustering potential was set to a value of 20 V. The data was evaluated manually using the Explorer tool of the SCIEX OS software, version 1.5.0.23389. For this purpose, a suitable time range of the eluting peak in the total ion current chromatogram (TIC) was summed to obtain a multiply charged spectrum. The 5 most intense multiple charge peaks were marked and deconvoluted using the Bio Tool Kit in the Explorer. The input spectrum isotope resolution was set to 1000 (very low). The mass step was 1 Da and the charge agent was set to H+. The mass range to be calculated was set between 170000 and 190000 Da.

##### Flow cytometric analysis of PROxAb Shuttle binding to CD33 cell surface receptors

Flow cytometric analyses were conducted at Reaction Biology Europe GmbH, Germany. For this, αCD33xMIC7 and αDIGxMIC7 were complexed with GNE987 (**6**) in different ratios resulting in different PROTAC-to-antibody-ratios as stated in the chapter “Generation of PROTAC-complexed PROxAb Shuttles”. Non-complexed PROxAb Shuttles and the control antibody, gemtuzumab with IgG4 PG-SPLE mutation (αCD33), were treated with the same procedure exchanging the DMSO dissolved PROTAC to DMSO only. Subsequently, the control antibody, complexed and non-complexed PROxAb Shuttles were stored at 4 °C overnight. On the next day, the control antibody, complexed and non-complexed PROxAb Shuttles were diluted in FACS-buffer (PBS 2% FCS 1 mM EDTA) at 1:100 (100 nM final antibody concentration, accordingly) and 1:1000 (10 nM final antibody concentration, accordingly) immediately before addition to cells. Cell lines (MV4-11, MOLM-13 and RAMOS) were pre-cultured for about 1 week in RPMI+Glutamax (Gibco) + 10% FCS (Bio&SELL) + 1% Penicillin/Streptomycin (Gibco). On the day of staining, cells were counted and plated at 2.0x10^5 c^ells per well in clear v-bottom 96-well microtiter plates (Thermofisher). Cells were spun down in the 96-well plate at 300xg for 5 min at 4 °C, supernatant was poured off, cell pellets were loosened by vortexing, and 100 µL PBS (Gibco) were added per well to wash cells. After centrifugation as before, PBS was poured off, and 100 µL of pre-diluted control antibody, complexed and non-complexed PROxAb Shuttles (1:100 and 1:1000) was added per well. Plates were sealed, vortexed, and staining incubation was performed for 1 h at 4 °C. Thereafter, 100 µL FACS-buffer was added per well, followed by centrifugation, pouring off of supernatant, vortexing and again addition of 100 µL FACS-buffer. After centrifugation as before, FACS-buffer was poured off, and 100 µL secondary antibody (goat anti-human FITC, Jackson, #109-095-098) was added per well at 1:100 dilution in FACS-buffer. Plates were sealed, vortexed, and staining incubation was performed for 1 h at 4 °C. Cells were washed as described above and finally resuspended in 100 µL FACS buffer prior to flow cytometric measurement using a CytoFLEX S flow cytometer (Beckman Coulter). 10.000 events were recorded per sample and subsequently analyzed using FlowJo software. For the analysis, mean fluorescence intensity (MFI) of the FITC-channel was assessed in single live cells, MFI of secondary antibody only samples were subtracted from all samples, duplicate samples were averaged and plotted using GraphPad prism.

##### Flow cytometric analysis of PROxAb shuttle internalization

Flow cytometric analyses were conducted at Reaction Biology Europe GmbH, Germany. The αCD33xMIC7 PROxAb variant was incubated for 2 h at 25 °C and 650 rpm in PBS with 5% VH032-pHAb dye (**15**) (dissolved in DMSO, Figure 3) at a 1.8-fold molar excess. The procedure was performed under the exclusion of light. CD33-positive MOLM13, MV4-11, U937 and receptor negative RAMOS cells were cultured in RPMI-1640 with 10% FCS. From each cell line, 200,000 cells were harvested, centrifuged, and incubated with either the dye-complexed PROxAb Shuttle or the dye-only control in 200 μL PBS with 1% FCS at a final concentration of 10 µg/mL. 6 h incubations were performed at 37 °C while shaking at 650 rpm in the dark. The cells were washed once with PBS containing 1% FCS and subsequently resuspended in 400 µL PBS with 1% FCS. Flowcytometric analyses were performed on a Becton Dickinson FACSCalibur flow cytometer. Staining of dead cells with propidium iodine (PI) was not performed to omit possible interference with VH032-pHAb dye (**15**). Quantitative analyses were done using the software FlowJo from BD Biosciences.

##### Assessment of BRD4 degradation via western blotting

Western Blotting was performed at Reaction Biology Europe GmbH, Germany. For degradation assays, CD33-positive MV4-11 cells were treated with GNE987-complexed αCD33 or αDIG PROxAb Shuttles at 1, 0.1, 0.01 and 0.001 nM. Unbound GNE987 (**6**) was used as a control at a concentration of 1 nM. MV4-11 cells were cultured in RPMI-1640 medium supplemented with 10% FCS and penicillin as well as streptomycin. MV4-11 cells were seeded in 12-well plates at 1 million cells/mL in 2 mL culture medium and cultured overnight at 37 °C and 5% CO_2_. PROxAb Shuttle complexation with PROTAC was performed as described previously. A control comprising unbound GNE987 (**6**) was preincubated at identical conditions, but in absence of a PROxAb Shuttle. Subsequently, cells were treated with either GNE987 (**6**)-complexed αCD33 or αDIG PROxAb shuttle at a PROTAC-to-antibody ratio of 1:1 or with GNE987 (**6**) using a Tecan D300e dispenser. All tested conditions were normalized to 0.0005% (v/v) DMSO and 3.x10^-5^ % (v/v) Tween-20. The cells were subsequently incubated at 37 °C and 5% CO_2_ for 24 h. Treated cells were harvested, washed in PBS and the resulting cell pellets were lysed in cell lysis buffer (20 mM TRIS pH 7.4, 100 mM NaCl, 1 mM EDTA, 0.5% TritonX‑100) supplemented with Roche cOmplete protease inhibitor mixture. After incubation for 10 min on ice, crude lysates were collected via centrifugation at 15,000xg, 4 °C and the resulting supernatants were precipitated by acetone. Dried pellets were dissolved in SDS loading buffer and the protein concentrations were determined with a Mettler Toledo UV5Nano photometer. Subsequently, samples (50 µg/lane) were applied to 4-12% Bis-Tris SDS-PAGE gels (Thermo Fisher Scientific). After completion of the SDS run, the samples were transferred onto nitrocellulose membranes (Sigma Aldrich) via western blotting. Next, membranes were blocked with 5% skim milk in 0.1% TBS-Tween-20 and incubated with primary antibodies against BRD4 (CST#13440, rabbit, 1:1000, Cell Signaling Technology) and Actin (PQ#679, mouse, 1:10,000, ThermoFisher) in 5% skim milk and 0.1% TBS-Tween-20. After incubation with corresponding HRP-labeled anti-rabbit/mouse secondary antibodies (GE‑Healthcare) at 1:10,000 dilution, the blots were developed with ECL solution (Advansta) and x-ray films (GE‑Healthcare). Afterwards, the obtained results were analyzed with the software ImageJ (version 1.53K, NIH). Background subtracted BRD4 signals were normalized to corresponding actin signals and the normalized BRD4 signal of the solvent control served as the reference point for the 100% value.

##### *In vitro* cytotoxicity assays

Cell viability assays were performed at Reaction Biology Europe GmbH, Germany. Therefore, cancer cells were seeded in different media into white cell culture-treated flat and clear bottom multi-well plates (corning^®^), followed by overnight incubation in a humid chamber at 37 °C and 5% CO_2_. PROxAb Shuttle complexation with PROTAC was conducted as described previously. Serial dilutions of PROxAb Shuttle solutions were added to the cells using nanodrop dispensing and a Tecan D300e Digital Dispenser. All wells were normalized to the same volume using solutions of 0.3% Tween-20 in PBS, pH 7.4 and DMSO to a final concentration of 0.05% DMSO and 0.003% Tween-20. Subsequently, treated cells were incubated at 37 °C and 5% CO_2_. The assays were developed after 3 days using CellTiter-Glo Luminescent Cell Viability Assay as described in the manufacturer’s protocol. In brief, the plates were equilibrated to room temperature for 30 minutes. Afterwards, 100 mL CellTiter-Glo Buffer were added to the CellTiter-Glo Substrate Flask and mixed well. Next, 25 µL of the reagent were transferred to each well of the cell culture plate. After incubation for 3 minutes at room temperature and shaking at 550 rpm, the plates were incubated for another 30 minutes at room temperature. The luminescence signals were measured on an Envision reader from Perkin Elmer. Solvent alone and staurosporine (at 10 µM) served as 100% viability and 0% viability controls, respectively. Raw data were converted into percent cell viability relative to solvent and staurosporine, which were set to 100% and 0% viability, respectively. IC50 value calculations were performed using GraphPad Prism software with a variable slope sigmoidal response fitting model.

##### *In vivo* pharmacokinetic studies

In preparation for the *in vivo* pharmacokinetic studies, αCD33 and αEGFR PROxAb Shuttles were complexed with the PROTAC GNE987 (**6**) or GNE987P (**7**) (αCD33 PROxAb Shuttle only) at a PROTAC-to-antibody ratio of 2:1, as described previously.

**αCD33 antibody, αCD33xMIC5, αCD33xMIC5 complexed with GNE987, αCD33xMIC7, αCD33xMIC7 complexed with GNE987**

Studies were performed in C57BL/6N inbred mice (n = 2 males and females for each group) that were provided by Charles River Laboratories Italia, Calco, Italy. The 7 – 8-week-old mice received 30 mg/kg (corresponding to 0.38 mg/kg PROTAC) of non-complexed and complexed αCD33xMIC5 and αCD33xMIC7 PROxAb Shuttles as well as αCD33 antibody as single doses that were intravenously injected into the tail vein. Blood samples were serially collected from all animals using a microsampling technique (20 μL for each blood withdrawal). After administration, 2 blood samples were taken on the first day and another 7 were taken during the following 3 weeks. Each sample was collected in a pre-chilled (0-4 °C) Minivette POCT EDTA tube, transferred in Microvette CB300 EDTA and centrifuged at 2500 x g for 10 min at 4 °C. The obtained plasma was transferred into a new vial and immediately stored at -80 °C until further analyses. The PK study as well as animal handling and experimentation were conducted in accordance with the Italian D.Lvo. 2014/26 and Directive 2010/63/EU. The study was performed at the Instituto di Ricerche Biomediche Antoine Marxer, Colleretto Giacosa, Italy. The institute is fully authorized by the Italian Ministry of Health. The total antibody concentration was determined by ligand binding assays (LBA) based on the Meso Scale Diagnostics technology (MSD, LLC., Rockville, MD). All incubation steps were performed at 22 °C with gentle agitation. All washing steps (200 µL/well) were performed with PBS-T, containing PBS, pH 7.4 and 0.01% Tween-20, using the plate washer ELx405 (BioTek instruments Inc., Winooski, VT). First, 2.5 µg/mL biotin-SP-conjugated AffiniPure goat anti-human IgG, Fcγ fragment specific (Jackson ImmunoResearch Europe Ltd., JIR, Cambridgeshire, United Kingdom, #109-065-098) was coated on MSD GOLD 96-well Streptavidin QUICKPLEX Plates (MSD, #L55SA) for 2 h. Afterwards, the plates were washed three times. Plasma samples, standards and quality controls were serially diluted in dilution buffer, consisting of PBS, pH 7.4, 0.05% Tween-20 and 3.0% (w/v) BSA, and incubated for 1 h. The plates were washed again and incubated for another hour with 0.6 µg/mL mouse anti-human IgG, F(ab’)2 fragment specific (JIR, #209-005-097), previously labeled with MSD GOLD SULFO-TAG (MSD, #R31AA-1) according to the manufacturer’s procedure. After a final washing step, 150 µL of 2x MSD Read Buffer T with surfactant (MSD, #R92TC) was added to each well and plates were read on a MESO Quickplex SQ120 plate reader (MSD). The Software Watson LIMS (Version 7.5, ThermoFisher Scientific Inc.) was used to fit the standard curve with a 5PL (Marquart) equation, weighting factor 1/Y^2^, and to calculate the total antibody (tAntibody) concentration of the plasma samples. The lower limit of quantification (LLOQ) of the method was 50 ng/mL. The concentration of GNE987 (**6**) was determined by liquid chromatography tandem mass spectrometry (LC-MS/MS) using a SCIEX 5500 triple quadrupole with Turbo Ion Spray source (ITS) in positive modality (SCIEX, Redwood City, CA, USA). Chromatographic separation was achieved using a Waters ACQUITY UPLC BEH (C18, 2.1 x 50 mm, 1.7 µm) column, mounted in a Waters ACQUITY I-class UPLC system (Milford, MA, USA), configured with a 100 µL extension loop. The chromatographic gradient used for phase A (H_2_O:acetonitrile 95:5, 0.1% formic acid) and B (acetonitrile:H_2_O 95:5, 0.1% formic acid) at a flow rate of 0.350 mL/min, was 0.25 min of 100% A isocratic, followed by a 2.25 min gradient to 100% B, with a subsequent 0.75 min of washing step at 100% B and 2.5 min of reconditioning at initial conditions. Extraction of GNE987 (**6**) from C57BL/6N mouse plasma samples was carried out by protein precipitation technique. In that manner, 3 µL of plasma sample were precipitated in 100 µL of acetonitrile containing 0.5 ng/mL of Exatecane-d5, used as internal standard, on a Phenomenex Impact Protein Precipitation Plate (Phenomenex, Torrance, CA, USA, CE0-7565). After 5 min of shaking at 900 rpm, all the wells were filtered by vacuum and collected in a clean 96-well plate. Subsequently, they were diluted with 100 µL of an aqueous solution containing 2.5% formic acid and submitted to LC-MS/MS analyses. All reagents were LC-MS grade or equivalent. The Software Watson LIMS (Version 7.5, ThermoFisher Scientific Inc.) was used to fit the standard curve on the area ratio (analyte signal/internal standard signal) on a linear regression, weighting factor 1/X^2^, and to calculate the total GNE987 PROTAC (**6**) concentration of the plasma samples. The lower limit of quantification (LLOQ) was 5 ng/mL, and the complete range of quantitation was 5-2000 ng/mL. The main pharmacokinetic parameters were estimated by noncompartmental analysis (NCA) using Phoenix WinNonlin version 8.3.4 (Pharsight Corporation, USA). The pharmacokinetic parameters have been obtained or calculated from the individual plasma concentrations of total antibody and GNE987 (**6**) analyte versus time after administration. Individual plasma concentration-time profiles were used for parameter estimation. The concentration of all pharmacokinetic samples that were calculated to be below the quantification limit (BQL) were considered as missing value to better estimate AUC_0-inf_, clearance and volume of distribution. The values for the terminal half-life (t_1/2_) and the first order rate constant associated with the terminal log-linear portion of the curve (ʎz) were calculated only when at least 3 time points were quantifiable in the terminal phase of the linear regression. Values below the quantification limit were considered 0 ng/mL for descriptive statistics.

**αCD33xMIC7+ GNE987P, αCD33xMIC7+ GNE987P administered sequentially and GNE987P alone**

The pharmacokinetic studies of αCD33xMIC7 complexed with GNE987P (**7**), αCD33xMIC7 and GNE987P (**7**) administered sequentially, as well as of GNE987P (**7**) alone were performed at EMD Serono Research & Development Institute, Inc. Billerica and are described in the following. All parts of the study plan concerning animal care have been reviewed by the EMD Serono Billerica site “Institutional Animal Care and Use Committee (IACUC)” which reports to the EMD Serono Billerica site Head. Protection of animals used, housing and welfare are guaranteed according to the Protocol number 20-002: Pharmacokinetic Studies in Rodents, Principal Investigator: Mary-Jo Miller. Physical facilities for accommodation and care of animals are in accordance with the provisions of the AAALAC (Association for Assessment and Accreditation of Laboratory Animal Care) and the EMD Serono animal facility is recorded as AAALAC unit #001473. For GNE987P (**7**) alone, female C57BL/6N inbred mice (composite profile, n=9) provided from Taconic (NY, USA) received a single tail vein intravenous (i.v.) bolus injection of 0.30 mg/kg in 2% (v/v) DMSO / 20% (v/v) (hydroxypropyl ß-cyclodextrin) Kleptose in water, at a dosing volume of 5 mL/kg. For the αCD33xMIC7 PROxAb Shuttle complexed with GNE987P (**7**), C57BL/6N inbred mice (n=12, composite profile) provided from Taconic (NY, USA) received a single tail vein intravenous (i.v.) bolus injection of 30 mg/kg of complexed αCD33xMIC7 PROxAb Shuttle (corresponding to 0.38 mg/kg PROTAC) in 5% (v/v) DMSO in PBS, at a dosing volume of 5 mL/kg. The last group was administered sequentially, first with the non-complexed αCD33xMIC7 PROxAb Shuttle at the same dose as in the group treated with the PROxAb Shuttle complex, followed 24h later by the PROTAC (GNE987P (**7**)) in the corresponding dose used for the αCD33xMIC7 PROxAb Shuttle complexed with GNE987P (**7**). For this, female C57BL/6N inbred mice (composite profile, n=12) provided from Taconic (NY, USA) received a single tail vein intravenous (i.v.) bolus injection of the αCD33xMIC7 PROxAb Shuttle of 30 mg/kg in 5% (v/v) DMSO in PBS, at a dosing volume of 5 mL/kg. The same mice, 24h after the administration of the PROxAb Shuttle, received a single tail vein intravenous (i.v.) bolus injection of GNE987P (**7**) at 0.40 mg/kg in 2% (v/v) DMSO/20% (v/v) (hydroxypropyl ß-cyclodextrin) Kleptose in water, at a dosing volume of 5 mL/kg. For the GNE987P (**7**) alone, consecutive blood samples (25 µL) were taken sub-lingually under light isoflurane anesthesia (n=3 per group (G)) at 0.1 (G1), 2 (G2), 6 (G3), 24 (G1), 72 (G2), 120 (G3), 240 (G2), 336 (G3) and 360h (G1) on K3-EDTA in pre-chilled Eppendorf tubes, stored on ice for at least 15 min before centrifugation at 2500 x g for 10 min at 4°C to obtain plasma. For the αCD33xMIC7 PROxAb Shuttle complexed with GNE987P (**7**) or the group administered sequentially with the αCD33xMIC7 PROxAb Shuttle followed by the PROTAC GNE987P (**7**), consecutive blood samples (25 µL) were taken sub-lingually under light isoflurane anesthesia (n=3 per group (G)) at 0.1 (G1), 6 (G2), 24 (G3), 96 (G4), 168 (G1), 240 (G2), 360 (G3), 600 (G4), 840 (G1), 960 (G2), 1080 (G3) and 1200h (G4) on K3-EDTA in pre-chilled Eppendorf tubes, stored on ice for at least 15 min before centrifugation at 2500 x g for 10 min at 4°C to obtain plasma. For the sample preparation, 10 µL plasma was diluted with 10 µL of methanol and precipitated with 80 µL of acetonitrile, containing labetalol as internal standard (2.5 µg/mL), in LowBind^®^ (protein) plates. After shaking/vortexing for 1 min, samples were filtered, (Captiva filtration on polypropylene filter, 0.45 µm pore size) and 120 µL of methanol:water (1:1, v/v) was added to the filtrate and stored at 4°C until analysis and put in the autosampler before injection. The analysis was carried out on a LC-MS/MS system consisting of an UPLC coupled to a QTRAP 6500+ (Sciex) mass spectrometer. Mobile phase A was water with 0.1% formic acid and mobile phase B was methanol with 0.1% formic acid. The gradient was started with 10% B to 95% B in 1.5 min and maintained at 95% B for 2 min, then decreased to 10% B in 0.5min and maintained to 10% B for 2 min. The chromatography was performed on a Poroshell 120 EC-C18 column, 2.7 µm particles, 3 x 50 mm, from Agilent Technologies. The flow rate was 0.6 mL/min and the cycle time (injection to injection) was approximately 6 minutes The sample injection volume was 10 µL. MRM transition for GNE987P (**7**) was 551.669 (m/z, z = 2) → 318.1 (m/z, z=2) and 329.101 (m/z, z=1) → 91 (m/z, z=1) for labetalol (IS). The calibration curve for quantitation was based on standards ranging from 0.5 (Lower Limit of Quantitation) to 10000 (Upper Limit of Quantitation) ng/mL, with 5 calibration points minimum and minimum 75% of calibration standards to be within ± 20% of their nominal values. The main pharmacokinetic parameters were estimated by noncompartmental analysis (NCA) using Phoenix WinNonlin version 8.3.4 (Pharsight Corporation, USA). The pharmacokinetic parameters have been calculated from the mean value of plasma concentrations of GNE987P (**7**) analyte versus time profile after administration. The concentration of all pharmacokinetic samples that were calculated to be below the lower limit of quantitation (LLOQ) were considered as missing value (i.e. values for time points at 24h and longer for GNE987P (**7**) alone, or 360h and longer for either the PROxAb Shuttle complexed with GNE987P (**7**) or for the sequential administration of PROxAb Shuttle followed by GNE987P (**7**)) to better estimate AUC_0-inf_, clearance and volume of distribution. The values for the terminal half-life (t_1/2_) and the first order rate constant associated with the terminal log-linear portion of the curve (λz) were calculated only when at least 3 time points were quantifiable in the terminal phase of the linear regression. Values below the LLOQ were considered 0 ng/mL for descriptive statistics.

**αEGFRxMIC7 complexed with GNE987 and GNE987 alone**

The pharmacokinetic studies of αEGFRxMIC7 complexed with GNE987 (**6**) and of GNE987 (**6**) alone were performed at EMD Serono Research & Development Institute, Inc. Billerica and are described in the following. All parts of the study plan concerning animal care have been reviewed by the EMD Serono Billerica site “Institutional Animal Care and Use Committee (IACUC)” which reports to the EMD Serono Billerica site Head. Protection of animals used, housing and welfare are guaranteed according to the Protocol number 20-002: Pharmacokinetic Studies in Rodents, Principal Investigator: Mary-Jo Miller. Physical facilities for accommodation and care of animals are in accordance with the provisions of the AAALAC (Association for Assessment and Accreditation of Laboratory Animal Care) and the EMD Serono animal facility is recorded as AAALAC unit #001473. For GNE987 (**6**) alone, female C57BL/6N inbred mice (composite profile, n=9) provided from Taconic (NY, USA) received a single tail vein intravenous (i.v.) bolus injection of 0.30 mg/kg in 2% (v/v) DMSO/20% (v/v) (hydroxypropyl ß-cyclodextrin) Kleptose in water, at a dosing volume of 5 mL/kg. For the αEGFRxMIC7 PROxAb Shuttle complexed with GNE987 (**6**), C57BL/6N inbred mice (composite profile, n=9) provided from Taconic (NY, USA) received a single tail vein intravenous (i.v.) bolus injection of 30 mg/kg of the αCD33xMIC7 PROxAb Shuttle (corresponding to 0.38 mg/kg PROTAC) in 5% (v/v) DMSO in PBS, at a dosing volume of 5 mL/kg. For the free GNE987 (**6**), consecutive blood samples (25 µL) were taken sub-lingually under light isoflurane anesthesia (n=3 per group (G)) at 0.1 (G1), 0.5 (G2), 1 (G3), 2 (G1), 4 (G3), 6 (G2), and 24h (G3) on K3-EDTA in pre-chilled Eppendorf tubes, stored on ice for at least 15 min before centrifugation at 2500 x g for 10 min at 4°C to obtain plasma. For the αEGFRxMIC7 PROxAb Shuttle complexed with GNE987 (**6**), consecutive blood samples (25 µL) were taken sub-lingually under light isoflurane anesthesia (n=3 per group (G)) at 0.1 (G1), 2 (G2), 6 (G3), 24 h (G1), 72 (G2), 168 (G3), 240 (G1), 432 (G2), 600 (G3), 720 (G1) and 960h (G2) on K3-EDTA in pre-chilled Eppendorf tubes, stored on ice for at least 15 min before centrifugation at 2500 x g for 10 min at 4°C to obtain plasma. The sample preparation, analysis of the PROTAC and the estimation of pharmacokinetic parameters were the same as described above for αCD33xMIC7+GNE987P (**7**) and GNE987P (**7**), except that the PROTAC analyzed here is GNE987 (**6**) instead of GNE987P (**7**). The MRM transition for GNE987 (**6**) was 548.788 (m/z, z = 2) → 779.2 (m/z, z=1) and 329.101 (m/z, z=1) → 91 (m/z, z=1) for labetalol (IS). Also, the concentrations for the 720 and 960 h time points after the administration of the αEGFRxMIC7 PROxAb Shuttle complexed with GNE987 (**6**) were calculated to be below the lower limit of quantitation (LLOQ) and were considered as missing value for the estimation of AUC_0-inf_, clearance and volume of distribution.

##### *In vivo* efficacy studies

*In vivo* efficacy studies encompassing αCD33 PROxAb Shuttles were performed at Nuvisan ICB, Germany. Therefore, 3 million MV4-11 cells were injected subcutaneously in the left flank of untreated female CB17 SCID mice. Randomization of the animals in the different treatment groups and the initiation of the treatment was started after the average tumor size reached 150 mm^3^. The control groups were treated with vehicle only (PBS pH 7.4, 5% DMSO (v:v)). The different groups were treated with 0.38 mg/kg GNE987 (**6**) and 30 mg/kg of either conjugated or unconjugated MIC7-based αCD33 PROxAb Shuttle (complexed with 0.38 mg/kg GNE987 (**6**)) once on day 1. Two additional groups were treated twice, either with 30 mg/kg of conjugated αCD33 PROxAb Shuttle (comprising 0.38 mg/kg GNE987 (**6**)) at day 1 followed by an additional treatment with 0.38 mg/kg GNE987 (**6**) at day 8 or with 0.38 mg/kg GNE987 at day 1 and 8. The individual groups were stopped before tumors reached a maximum tumor volume (1600 mm^3^).

**Supplementary Figures and Tables**

#### Figures

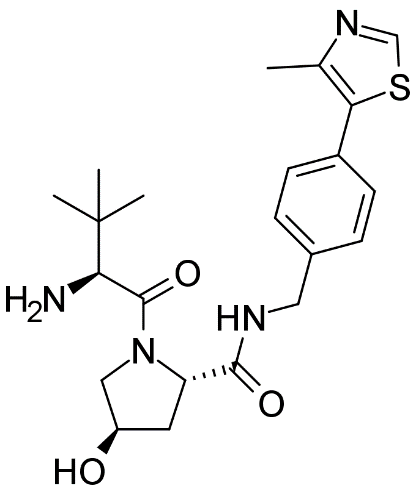

**Figure S1.** Structure of VHL Ligand VH032 (**5**).

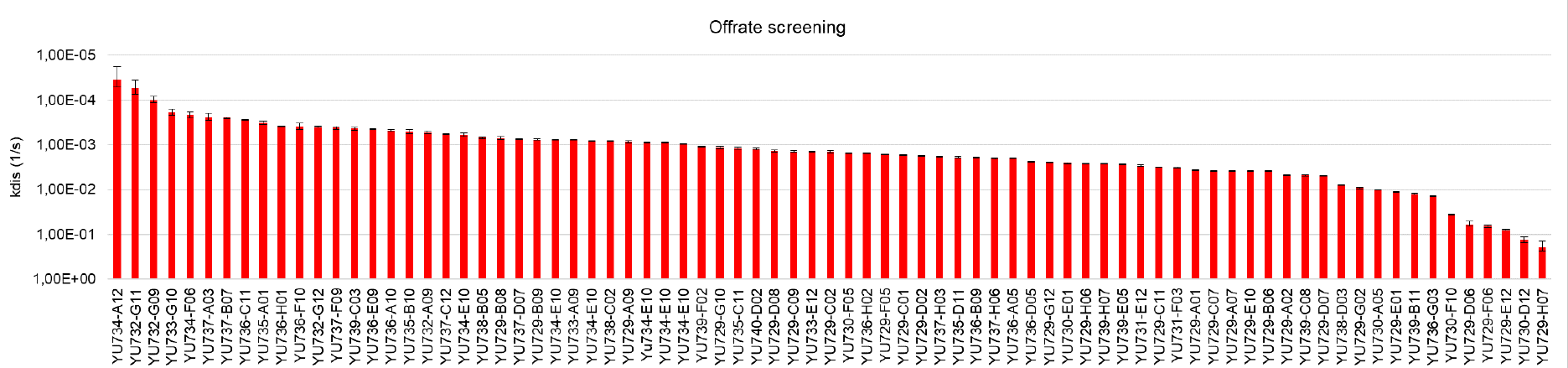

**Figure S2.** Off-rate screening using Bio-Layer Interferometry (BLI).The clones YU733-G10 and YU734-F06 refer to MIC5 and MIC7, respectively.

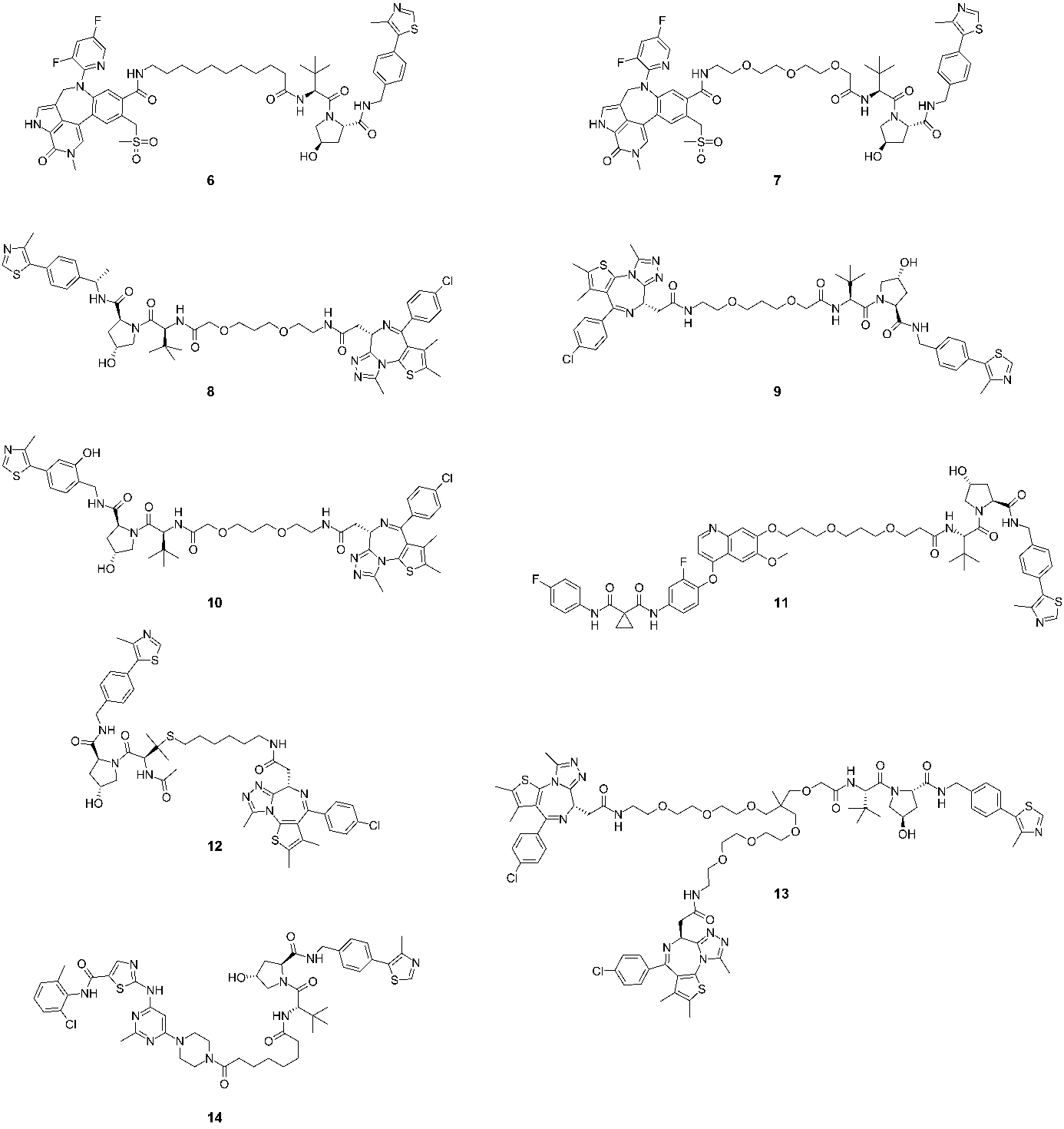

**Figure S3.** Structures of PROTACs used for SPR analytics: **6** (GNE987), **7** (GNE987P), **8** (ARV771), **9** (BETTY2), **10** (BETTY3), **11** (cMETd1); **12** (AT1), **13** (SIM1), and **14** (SIAIS178)

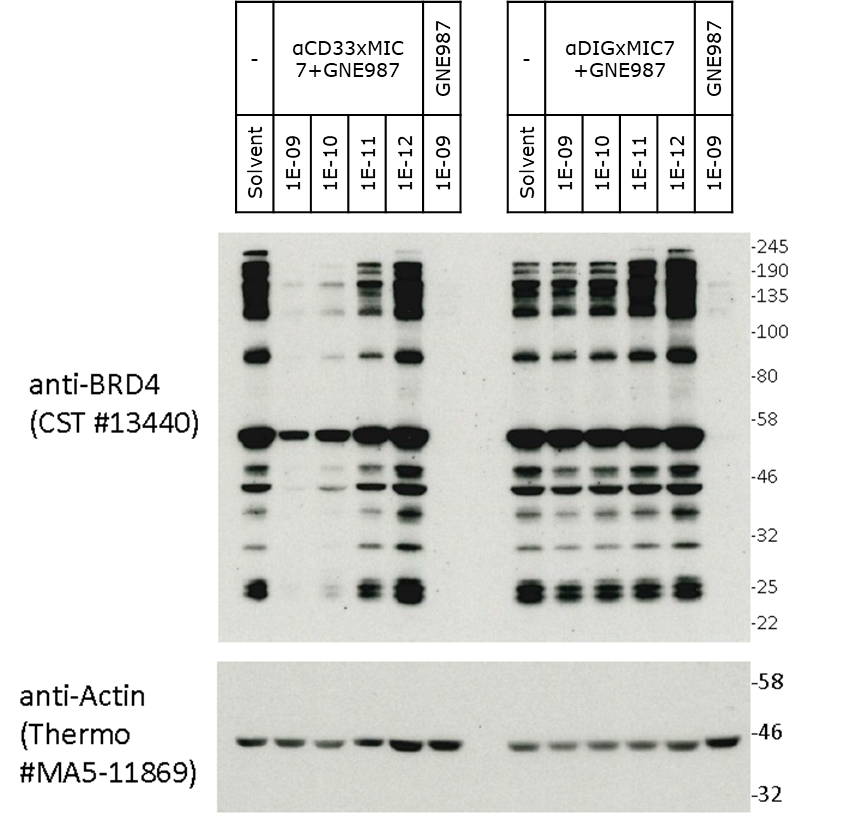

**Figure S4.** Western blot of MV4-11 cells treated with αCD33xMIC7 complexed with PROTAC GNE987 (**6**), negative control αDIGxMIC7+GNE987 (**6**) in serial dilution and PROTAC GNE987 (**6**) alone at 1 nM. All PROxAb Shuttles comprised a PROTAC-to-antibody ratio of 1:1.

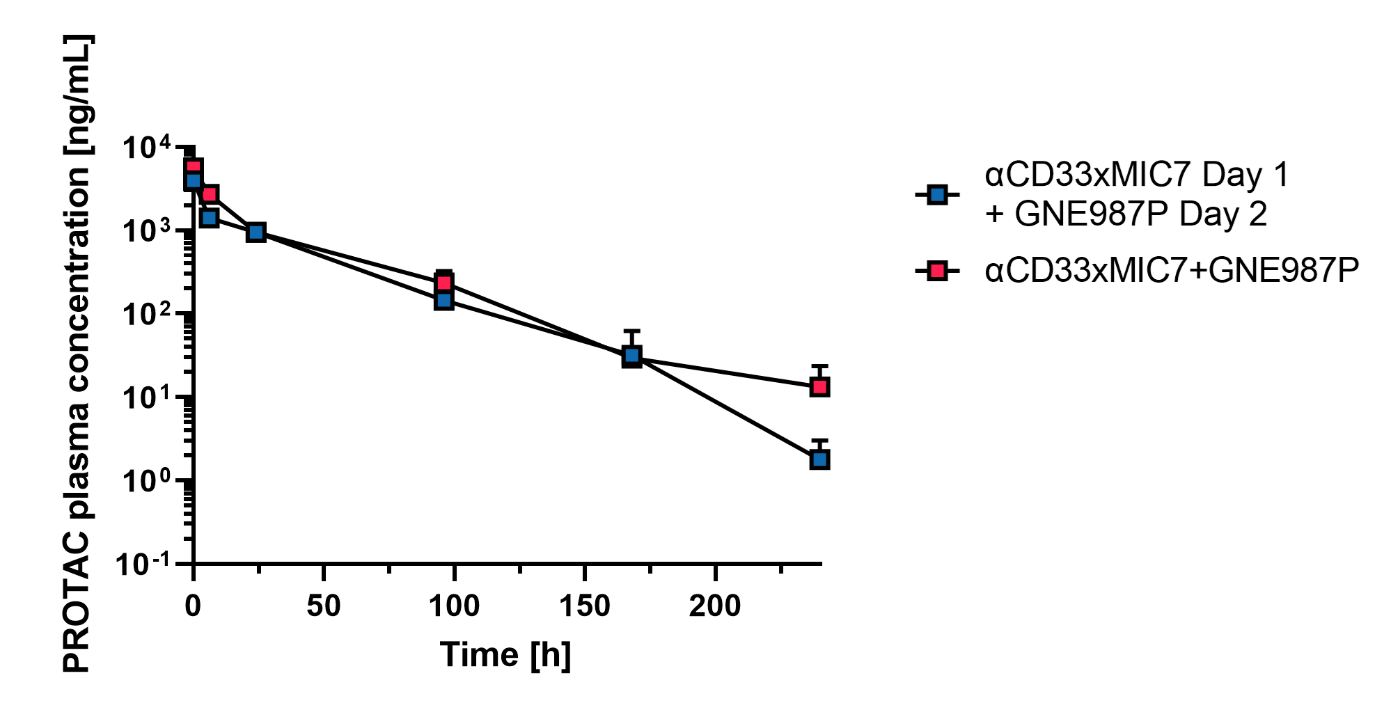

**Figure S5**. Pharmacokinetic analyses of total PROTAC. Total PROTAC profiles after administration of αCD33xMIC7 Shuttle after complexation with GNE987P (**7**) (red squares) and of sequentially administered non-complexed αCD33xMIC7 PROxAb Shuttle followed by administration of GNE987P (**7**) (blue squares) 24 hours later.

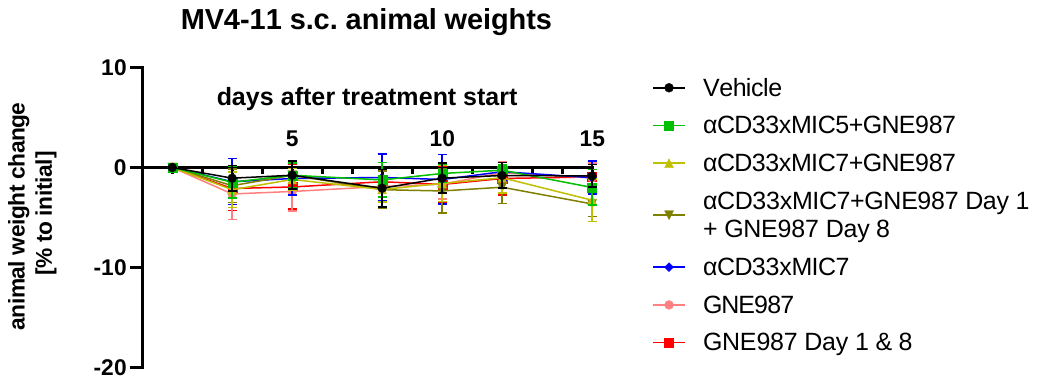

**Figure S6**. Animal Weights of MV4-11 tumor-bearing mice over time after indicated treatment.

#### Tables

**Table S1.** Affinity parameters of αCD33xMIC7 and αEGFRxMIC7 to different PROTACs from SPR measurements at 25 °C. N/D: not determined.

|  | **αCD33xMIC7** | | | **αEGFRxMIC7** | | |
| --- | --- | --- | --- | --- | --- | --- |
| **PROTAC** | **KD [M]** | **k_on_ [1/Ms]** | **k_off_ [1/s]** | **K_D_ [M]** | **k_on_ [1/Ms]** | **k_off_ [1/s]** |
| GNE987 (**6**) | 4.19E-10 | 2.49E+05 | 1.01E-04 | <1E-09 | N/D | <1.00E-4 |
| GNE987P (**7**) | 1.96E-09 | 4.97E+05 | 9.23E-04 | <1E-09 | 6.53E+05 | 3.20E-04 |
| ARV771 (**8**) | 1.07E-08 | 7.88E+05 | 3.92E-03 | 5.39E-09 | 1.39E+05 | 7.48E-04 |
| BETTY2 (**9**) | 2.11E-09 | 3.10E+05 | 6.50E-04 | 1.06E-09 | 3.15E+05 | 3.25E-04 |
| BETTY3 (**10**) | 3.26E-09 | 7.12E+05 | 2.31E-03 | 1.86E-09 | 3.77E+05 | 6.93E-04 |
| cMETd1 (**11**) | 1.23E-08 | 1.85E+04 | 2.28E-04 | 9.37E-09 | 1.85E+04 | 1.72E-04 |
| AT1 (**12**) | 2.07E-09 | 1.08E+06 | 2.23E-03 | 1.20E-09 | 2.95E+05 | 3.55E-04 |
| SIM1 (**13**) | 4.76E-09 | 1.54E+05 | 9.27E-04 | 1.90E-09 | 6.71E+04 | 1.27E-04 |
| SIAIS178 (**14**) | 8.31E-09 | 1.14E+04 | 9.50E-05 | <1E-09 | N/D | <1.00E-4 |

**Table S2**. Potencies of PROxAb Shuttles targeting different tumor associated antigens (TAA) (αCD33xMIC7 or αEGFRxMIC7, respectively) complexed with GNE987 (**6**) or GNE987P (**7**) (PROTAC-to-antibody ratio of 1:1) and free PROTAC GNE987 (**6**) or GNE987P (**7**) on CD33-positive and EGFR-positive cells as well as respective negative cell lines (RAMOS for CD33 and HEPG2 for EGFR). N/T: not tested.

| **Cell line** | **TAA** | **αTAAxMIC7 + GNE987** | **αDIGxMIC7+ GNE987** | **GNE987 (6)** | **αTAAxMIC7+ GNE987P** | **αDIGxMIC7+ GNE987P** | **GNE987P** |
| --- | --- | --- | --- | --- | --- | --- | --- |
| MV411 | CD33 | 0.28 nM | 8.1 nM | < 0.03 nM | 0.059 nM | 35 nM | 0.26 nM |
| MOLM13 | CD33 | 8.1 nM | >100 nM | 0.020 nM | 6.9 nM | >100 nM | 3.7 nM |
| RAMOS | CD33 | >100 nM | >100 nM | 0.089 nM | >100 nM | >100 nM | 15 nM |
| A431 | EGFR | 0.26 nM | >100 nM | 0.25 nM | N/T | N/T | N/T |
| MDAMB468 | EGFR | 0.27 nM | >100 nM | 0.16 nM | N/T | N/T | N/T |
| HEPG2 | EGFR | >100 nM | >100 nM | 2.2 nM | N/T | N/T | N/T |

**Table S3.** IC50 values of PROxAb Shuttles targeting various tumor-associated antigens such as HER2, BCMA, NAPI2B and EGFR complexed with PROTACs GNE987 (**6**), GNE987P (**7**) or SIM1 (**13**) on receptor-positive cell lines. The PROTAC-to-antibody ratio (PAR) was adjusted according to the PAR column.

| **Cell line** | **Target** | **PROxAb Shuttle clone** | **PROTAC** | **IC50 [M] PROTAC** | **IC50 [M]**  **αTAAxMIC7** | **IC50 [M]**  **αDIGxMIC7** | **PAR** | **Antibody Isotype** |
| --- | --- | --- | --- | --- | --- | --- | --- | --- |
| **BT474** | HER2 | MIC7 | GNE987 | 7.5E-11 | 1.5E-10 | 1.7E-08 | 1.0 | IgG1-PG-LALA |
| **BT474** | HER2 | MIC7 | GNE987P | 2.6E-09 | 2.0E-10 | 3.5E-08 | 1.0 | IgG1-PG-LALA |
| **MDAMB453** | HER2 | MIC7 | GNE987 | 9.4E-11 | 8.7E-11 | 4.5E-08 | 1.0 | IgG1-PG-LALA |
| **MDAMB453** | HER2 | MIC7 | GNE987P | 4.4E-09 | 5.2E-10 | 3.0E-08 | 1.0 | IgG1-PG-LALA |
| **SKBR3** | HER2 | MIC7 | GNE987 | 1.5E-10 | 1.9E-10 | 2.1E-09 | 1.0 | IgG1-PG-LALA |
| **SKBR3** | HER2 | MIC7 | GNE987P | 1.5E-08 | 2.1E-10 | 5.3E-08 | 1.0 | IgG1-PG-LALA |
| **K562** | BCMA | MIC7 | GNE987 | 2.0E-10 | 1.9E-07 | 2.9E-08 | 1.0 | IgG1-PG-LALA |
| **K562** | BCMA | MIC7 | GNE987P | 2.8E-08 | > 1.0E-07 | > 1.0E-07 | 1.0 | IgG1-PG-LALA |
| **LP1** | BCMA | MIC7 | GNE987 | 7.8E-11 | 3.4E-08 | > 1.0E-07 | 1.0 | IgG1-PG-LALA |
| **LP1** | BCMA | MIC7 | GNE987P | 4.7E-09 | 3.4E-08 | > 1.0E-07 | 1.0 | IgG1-PG-LALA |
| **NCIH929** | BCMA | MIC7 | GNE987 | 1.3E-11 | 2.0E-10 | 3.2E-08 | 1.0 | IgG1-PG-LALA |
| **NCIH929** | BCMA | MIC7 | GNE987P | 5.6E-10 | 9.1E-11 | 2.8E-08 | 1.0 | IgG1-PG-LALA |
| **OPM2** | BCMA | MIC7 | GNE987 | < 3.0E-11 | 3.6E-09 | 5.1E-08 | 1.0 | IgG1-PG-LALA |
| **OPM2** | BCMA | MIC7 | GNE987P | 4.8E-10 | 9.9E-10 | 3.8E-08 | 1.0 | IgG1-PG-LALA |
| **OVCAR3** | NAPi2B | MIC7 | GNE987 | 2.9E-10 | 2.8E-10 | > 1.0E-07 | 1.0 | IgG1-PG-LALA |
| **MOLM13** | B7H3 | MIC7 | GNE987 | 1.9E-10 | 4.0E-10 | 4.6E-08 | 1.5 | IgG4-PG-SPLE |
| **MOLM13** | B7H3 | MIC7 | GNE987P | 5.8E-09 | 1.5E-07 | N/T | 1.5 | IgG4-PG-SPLE |
| **MV411** | B7H3 | MIC7 | GNE987 | 4.7E-11 | 9.0E-11 | 1.2E-08 | 1.5 | IgG4-PG-SPLE |
| **MV411** | B7H3 | MIC7 | GNE987P | 4.0E-10 | 2.4E-09 | N/T | 1.5 | IgG4-PG-SPLE |
| **U937** | B7H3 | MIC7 | GNE987 | 2.3E-10 | 3.9E-10 | 1.0E-07 | 1.5 | IgG4-PG-SPLE |
| **U937** | B7H3 | MIC7 | GNE987P | 1.5E-09 | 3.2E-08 | N/T | 1.5 | IgG4-PG-SPLE |
| **MOLM13** | CLL1 | MIC7 | GNE987 | 5.4E-11 | 4.4E-08 | > 1.0E-07 | 1.5 | IgG4-PG-SPLE |
| **MOLM13** | CLL1 | MIC7 | GNE987P | 3.6E-09 | 3.5E-08 | > 1.0E-07 | 1.5 | IgG4-PG-SPLE |
| **U937** | CLL1 | MIC7 | GNE987 | 1.2E-10 | 6.0E-08 | > 1.0E-07 | 1.5 | IgG4-PG-SPLE |
| **U937** | CLL1 | MIC7 | GNE987P | 2.1E-09 | 2.3E-08 | > 1.0E-07 | 1.5 | IgG4-PG-SPLE |
| **K562** | CLL1-negative | MIC7 | GNE987 | 3.7E-10 | > 1.0E-07 | > 1.0E-07 | 1.5 | IgG4-PG-SPLE |
| **K562** | CLL1-negative | MIC7 | GNE987P | 4.2E-08 | > 1.0E-07 | > 1.0E-07 | 1.5 | IgG4-PG-SPLE |
| **A431** | EGFR | MIC7 | GNE987P | 1.6E-07 | 5.6E-10 | > 1.0E-07 | 1.0 | IgG1-PG-LALA |
| **A431** | EGFR | MIC7 | SIM1 | 3.5E-09 | 5.5E-10 | > 1.0E-07 | 1.0 | IgG1-PG-LALA |
| **MDAMB468** | EGFR | MIC7 | GNE987P | 4.6E-08 | 6.8E-10 | > 1.0E-07 | 1.0 | IgG1-PG-LALA |
| **MDAMB468** | EGFR | MIC7 | SIM1 | 5.7E-09 | 8.1E-10 | > 1.0E-07 | 1.0 | IgG1-PG-LALA |
| **MV411** | CD33 | MIC5 | GNE987P | 5.2E-10 | 1.3E-10 | - | 0.5 | IgG4-PG-SPLE |
| **MV411** | CD33 | MIC5 | GNE987P | 5.2E-10 | 1.1E-10 | - | 1.0 | IgG4-PG-SPLE |
| **MV411** | CD33 | MIC5 | GNE987P | 5.2E-10 | 8.2E-12 | - | 1.5 | IgG4-PG-SPLE |
| **MV411** | CD33 | MIC5 | GNE987 | 9.0E-12 | 3.2E-11 | - | 1.0 | IgG4-PG-SPLE |
| **MOLM13** | CD33 | MIC5 | GNE987 | 4.7E-11 | 4.4E-09 | - | 1.0 | IgG4-PG-SPLE |

**Table S4.** SPR affinity data for αCD33xMIC5 and αCD33xMIC7 binding to GNE987 (**6**) or GNE987P (**7**) at 37 °C.

| PROTAC | αCD33xMIC5 | | | αCD33xMIC7 | | |
| --- | --- | --- | --- | --- | --- | --- |
|  | **k_on_ (1/Ms)** | **k_off_ (1/s)** | **K_D_ (M)** | **k_on_ (1/Ms)** | **k_off_ (1/s)** | **K_D_ (M)** |
| 6 (GNE987) | 3.84E+05 | 2.19E-03 | 5.70E-09 | 2.49E+05 | 1.04E-04 | 4.19E-10 |
| 7 (GNE987P) | - | - | - | 2.70E+05 | 8.1E-04 | 2.98E-09 |

**Table S5.** Affinities (K_D_), change in enthalpy (ΔH) and stoichiometry (N) of wildtype αCD33xMIC7 and humanized variants αCD33xMIC7_1.1-MIC7_1.6 when binding to VH032 derived from isothermal calorimetry measurements.

| **Clone** | **Backbone** | **K_D_ [nM]** | **ΔH [kJ mol^-1^]** | **N** |
| --- | --- | --- | --- | --- |
| **αCD33xhMIC7_1.6** | IgG1 PG-LALA | <1 | -71 | 2.0 |
| **αCD33xhMIC7_1.5** | IgG1 PG-LALA | <1 | -80 | 2.0 |
| **αCD33xhMIC7_1.4** | IgG1 PG-LALA | ~3 | -69 | 2.1 |
| **αCD33xhMIC7_1.3** | IgG1 PG-LALA | <1 | -69 | 2.0 |
| **αCD33xhMIC7_1.2** | IgG1 PG-LALA | <2 | -61 | 2.0 |
| **αCD33xhMIC7_1.1** | IgG1 PG-LALA | <1 | -49 | 2.1 |
| **αCD33xMIC7_1.0** | IgG1 PG-LALA | <1 | -57 | 1.8 |
| **αCD33xMIC7** | IgG4 PG-SPLE | 5 | -52 | 2,1 |

**Table S6.** Affinity parameters of humanized αCD33xhMIC7_1.6 binding to PROTAC GNE987 (**6**) and GNE987P (**7**) derived from SPR measurements at 25 °C.

| **Name** | **k_on_ [1/M]** | **k_off_ [1/s]** | **K_D_ [1/Ms]** |
| --- | --- | --- | --- |
| **6 (**GNE987) | 3.2E+04 | 3.9E-05 | 1.2E-09 |
| **7** (GNE987P) | 2.6E+05 | 2.2E-04 | 8.6E-10 |

#### Ethical statement

*In vivo* studies were either performed at Istituto di Ricerche Biomediche “Antoine Marxer” – RBM Colleretto Giacosa (TO), Italy, Nuvisan ICB GmbH, Berlin, Germany or EMD Serono Research & Development Institute, Inc. Billerica, MA 01821, USA. At at Istituto di Ricerche Biomediche “Antoine Marxer” – RBM Colleretto Giacosa all parts of the studies concerning animal care have been either reviewed by the RBM Designed Veterinarian and Animal Welfare Officer. Protection of animals used, housing and welfare are guaranteed according to the Italian D.Lvo No. 26 of March 4, 2014. Physical facilities for accommodation and care of animals are in accordance with the provisions of the Italian D.Lvo 2014/26 and of Directive 2010/63/EU. The Institute is fully authorized by Italian Ministry of Health. For studies performed at Nuvisan ICB, Berlin all animal experiments were approved by the local ethics committee on animal research (Landesamt für Gesundheit und Soziales Berlin, Bereich des Veterinärwesens, Germany, LaGeSo No. G 0200/19). At EMD Serono Research & Development Institute, Inc. Billerica all parts of the study plan concerning animal care have been reviewed by the EMD Serono Billerica site “Institutional Animal Care and Use Committee (IACUC)” which reports to the EMD Serono Billerica site Head. Protection of animals used, housing and welfare are guaranteed according to the Protocol number 20-002: Pharmacokinetic Studies in Rodents, Principal Investigator: Mary-Jo Miller. Physical facilities for accommodation and care of animals are in accordance with the provisions of the AAALAC (Association for Assessment and Accreditation of Laboratory Animal Care) and the EMD Serono animal facility is recorded as AAALAC unit #001473.

Livestock farming and immunization of New World Camelids (llamas, alpacas, huarizos) was managed by preclinics GmbH according to the relevant national and international guidelines. The study was approved by an Institutional Animal Care and Use Committee, the Lower Saxony State Office for consumer protection and food safety (Oldenburg, Germany, reference number 33.19-42502-05-17A210).
